# A Serum- and Feeder-Free System to Generate CD4 and Regulatory T Cells from Human iPSCs

**DOI:** 10.1101/2023.07.01.547333

**Authors:** Helen Fong, Matthew Mendel, John Jascur, Laeya Najmi, Ken Kim, Garrett Lew, Swetha Garimalla, Suruchi Schock, Jing Hu, Andres Villegas, Anthony Conway, Jason D. Fontenot, Simona Zompi

**Affiliations:** Sangamo Therapeutics, 501 Canal Blvd. Richmond, CA 94804; Proximity Therapeutics, 135 Mississippi Street, San Francisco, CA 94107; BioMarin, 105 Digital Dr, Novato, CA 94949; OmniAb, 5980 Horton Street, Suite 600 Emeryville, CA 9460; Radar Therapeutics, 2625 Durant Ave, Berkeley, CA 94720; Replay, 5555 Oberlin Drive, Unit 120, San Diego, CA 92121; Altos Labs, 1300 Island Drive, Redwood City, CA 94065

## Abstract

iPSCs can serve as a renewable source of a consistent edited cell product, overcoming limitations of primary cells. While feeder-free generation of clinical grade iPSC-derived CD8 T cells has been achieved, differentiation of iPSC-derived CD4sp and regulatory T cells requires mouse stromal cells in an artificial thymic organoid. Here we report a serum- and feeder-free differentiation process suitable for large-scale production. Using an optimized concentration of PMA/Ionomycin, we generated iPSC-CD4sp T cells at high efficiency and converted them to Tregs using TGFβ and ATRA. Using genetic engineering, we demonstrated high, non-viral, targeted integration of an HLA-A2 CAR in iPSCs. iPSC-Tregs +/- HLA-A2-targeted CAR phenotypically, transcriptionally and functionally resemble primary Tregs and suppress T cell proliferation *in vitro*. Our work is the first to demonstrate an iPSC-based platform amenable to manufacturing CD4 T cells to complement iPSC-CD8 oncology products and functional iPSC-Tregs to deliver Treg cell therapies at scale.

## Introduction

Engineered autologous T cell therapies have shown the ability to transform the oncology field, yet producing these products at scale for large numbers of patients has proven challenging. The ability to derive hypoimmunogenic engineered T cells from self-renewing human induced pluripotent stem cells (iPSCs) could significantly increase access to these therapies^1–3^. The advantages of using iPSCs over primary cells include the capacity to easily multiplex edit the starting cell source, access to a self-renewable and well-characterized product, and standardization of the product through the development of a master cell bank (MCB). However, while using iPSCs as source material can have numerous benefits in terms of standardization and scale-up, differentiation of T cells from iPSCs has been challenging.

Access to a self-renewable, potentially unlimited source of cells is particularly attractive for cell types that are difficult to isolate or present in small numbers, such as regulatory T cells (Tregs). In recent years there has been much interest in the use of Tregs to treat a wide variety of diseases, including autoimmune disorders, as well as in organ transplantation^4^. As of 2022, there were more than 30 active and completed clinical trials evaluating the safety and efficacy of Treg cell therapies^5^. Most of the studies have focused on *in vitro*-expanded polyclonal Tregs but showing limited clinical efficacy. Promising preclinical results testing antigen-specific CAR Tregs^6^ led to the first CAR Treg clinical trial authorization (NCT04817774) for kidney transplant patients, using *ex vivo,* autologous, naïve Tregs engineered with a CAR construct recognizing HLA-A2, to investigate the induction of tolerance in HLA-A2 negative patients receiving an HLA-A2 positive renal graft.

Currently, sourcing CD4^+^ T cells and Tregs for cell therapies is invasive, time-consuming, and yields small numbers, particularly of Tregs which comprise only 5-7% of the circulating CD4 T cell population^7^. Furthermore, CD4^+^ T cells and Tregs obtained from these sources are polyclonal in nature and can introduce variability and non-specificity. While we have been successful in introducing a CAR construct in primary Tregs to mediate a specific immunosuppressive response^6^, a process for generating specific CD4^+^ T cells and Tregs, especially genetically engineered cells, with high efficiency and at clinically relevant scale has remained a challenge.

Numerous studies have shown the ability to generate large quantities of T cells from iPSCs, focusing primarily on the differentiation to cytotoxic T lymphocytes (CTLs)^8–10^. Improvements have recently been made in the differentiation processes to generate iPSC-derived CAR CD8 single positive (CD8sp) T cells that more closely resemble primary CD8αβ^+^ T cells rather than innate γδ T cells, by knocking out *TRAC* and delaying and attenuating CAR expression^11, 12^. The artificial thymic organoid (ATO) system has also proven to be successful in generating functional CD3^+^CD8αβ^+^ T cells that produce IFN-γ, TNF-α, and IL-2 and proliferate upon stimulation^10^. Recent data from the Human Cell Atlas group reported that these CD8 T cells differentiated *in vitro* were most similar to type 1 innate T cells - a prenatal unconventional T cell type particularly abundant during early development in the fetus that expresses high levels of *ZBTB16* (Promyelocytic Leukemia Zinc Finger Protein or PLZF) and has an innate-like transcriptional profile^13^. The authors further hypothesize that unconventional T cells arise *via* thymocyte-thymocyte (T-T) interactions during development as well as in the ATO system which does not contain any thymic epithelial cells (TECs)^13^.

Interestingly, the ATO system also produces a small population of CD3^+^CD4^+^ T cells^10, 14^ that can be converted to FOXP3^+^ Tregs^15^. However, a scalable method to generate clinically relevant numbers of CD4^+^ T cells and Tregs from iPSCs has yet to be demonstrated. This need has become more compelling in recent years as CD4^+^ T cells are known to enhance CD8^+^ CTL responses^16^ and have been shown to be therapeutically useful in CAR-T cell therapies, increasing product potency^17, 18^ and demonstrating long-term functional persistency in clinical trials^19^.

Current methods to generate CD8sp T cells, CD4 single positive (CD4sp) T cells and Tregs from iPSCs utilize serum or mouse stromal cells at some stage of the differentiation process making it difficult for use in clinical development. In addition, the use of an ATO system is challenging to scale-up for manufacturing as it requires the maintenance of a 3D system and involves dissociation and extraction of the cells of interest from a mixed population of mouse and human cells. Here we show for the first time the ability to efficiently generate both CD4sp T cells and Tregs from iPSCs in completely serum- and feeder-free conditions. We identified a potent cell stimulator to convert CD4^+^CD8^+^ double positive T (DP T) cells to CD4sp T cells at high efficiency that can be further converted to FOXP3^+^ Tregs using TGFβ, all-trans retinoic acid (ATRA), and IL-2 with CD3/CD28/CD2 activation. We show that the iPSC-CD4sp T cells have an innate-like transcriptional profile, including *ZBTB16* expression, evocative of prenatal unconventional T cells^13^, can be activated upon stimulation and secrete IL-2 and IL-4. The iPSC-Tregs possess typical Treg characteristics, are transcriptionally more similar to primary Tregs than primary conventional T (Tconv) cells and suppress primary CD4 Tconv proliferation. In addition, iPSCs engineered to express an HLA-A2 CAR and a fully functional TCR under the control of the *TRAC* promoter could effectively differentiate to CD4sp T cells and Tregs that suppress Tconv proliferation *via* activation through the TCR or the HLA-A2 CAR. Finally, we show that the iPSC-Tregs are amenable to cryopreservation and retain function post-thaw. This novel method provides a viable path forward towards the development of a scalable iPSC-derived Treg cell therapy for autoimmune diseases. This novel process provides a high yield of innate-like iPSC-CD4sp cells. Further optimization of culture media composition and/or cell engineering approaches should achieve a fully mature iPSC-derived CD4sp T cells which could enhance the iPSC-derived CD8sp T cell products currently in the clinic for oncology applications^17, 19, 20^.

## Results

### Generation of CD34^+^ hematopoietic stem and progenitor cells (HSPCs) and T cell progenitors from iPSCs

iPSCs were derived from sorted CD4^+^CD25^+^ cells from a healthy human donor. Derivation of iPSCs from T cells has been previously documented with resulting T cells differentiated from reprogrammed iPSCs possessing antigen specificity due to retention of a rearranged TCR^9, 21, 22^. Additionally, evidence suggests improved T cell lineage differentiation when iPSCs originate from mature T cells due to removal of the barrier to TCR rearrangement^9, 23^. CD4^+^CD25^+^ T cells were isolated and FACS-purified from healthy human leukapheresis blood and reprogrammed by Sendai virus carrying vectors encoding for the four Yamanaka factors: OCT-4, SOX2, KLF4, and C-MYC. Multiple clones were generated and characterized by immunocytochemistry to confirm expression of OCT4, SOX2, TRA-1-81, and SSEA-4 **(Supp. Figure 1A)**. Pluripotency was confirmed using the PluriTest assay (ThermoFisher, **Supp. Figure 1B**). In addition, G-band karyotyping confirmed the absence of any chromosomal aberrations **(Supp. Figure 1C)**.

To facilitate future translation from discovery to process development we utilized serum-free, commercially available reagents from the iPSC stage to the DP T stage. As a first step, a modified version of the spin embryoid body (EB) method was used to differentiate the iPSCs into early stages of hematopoiesis from days 0 to 8, followed by expansion of the hematopoietic stem and progenitor cells from days 8 to 14^24, 25^ **(Figure 1A)**. The EBs increased in size over time and by day 14, numerous free-floating HSPCs emerged around the periphery expressing both CD34 and CD45 (65%±13.5%, n=3) **(Figure 1B and 1C)**. The approximate yield of HSPCs per input iPSC was 16.5±9.3, n=3 **(Figure 2)**.

**Figure 1.**
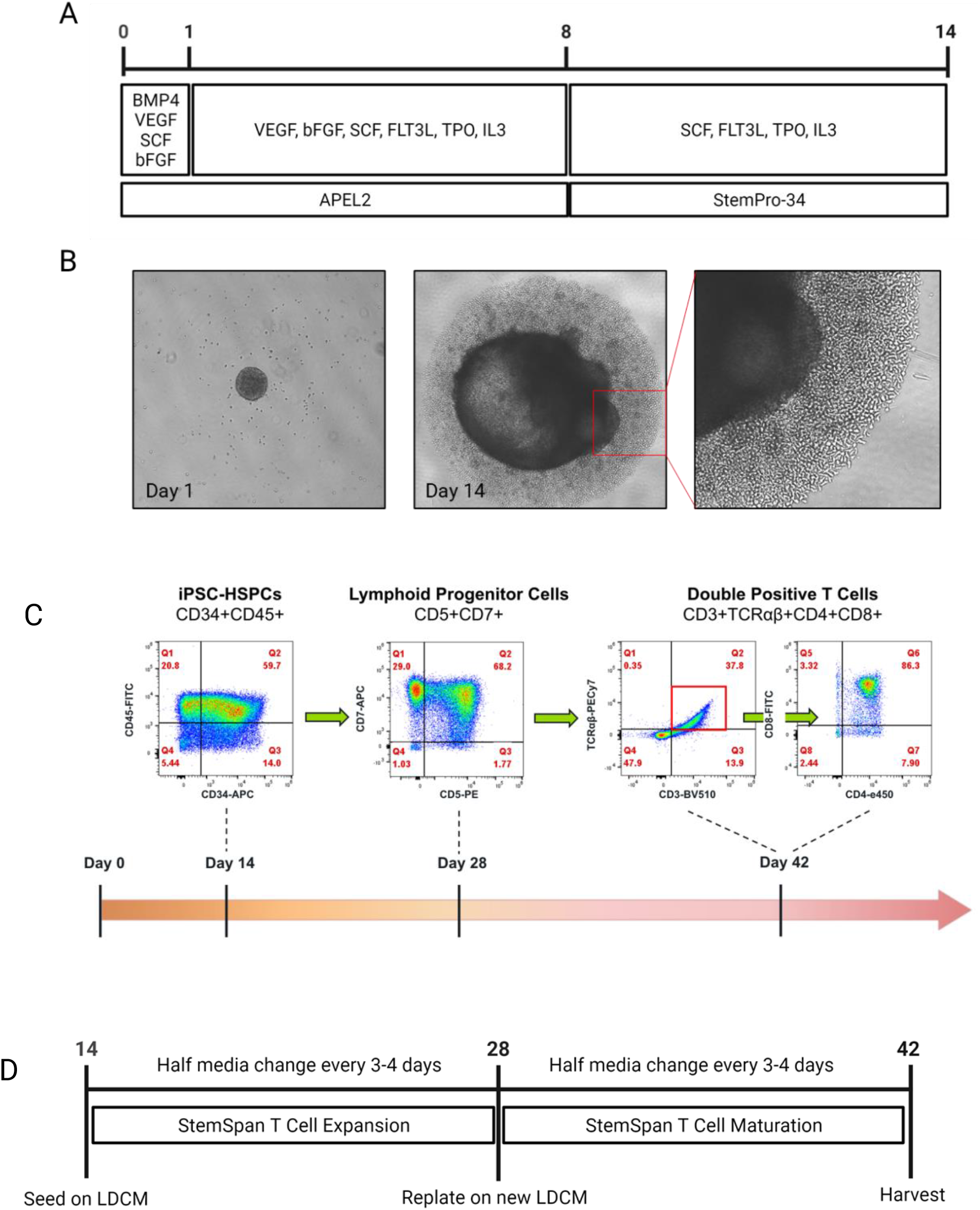
Differentiation of iPSCs to hematopoietic stem and progenitor cells (HSPCs), lymphoid progenitors and double positive (DP) T cells in serum- and feeder-free conditions. A. Schematic for differentiation of iPSCs to HPSCs. B. Embryoid bodies (EBs) at day 1 and 14 of differentiation. HSPCs emerged from the EB by day 14 and appeared as loosely packed cells (inset). C. Representative flow cytometric analyses of HSPCs, lymphoid progenitor cells, and DP T cells from iPSCs. Cells were analyzed for phenotype over a 42-day period at the indicated timepoints. Data show one representative flow plot set of n=3 independent experiments from n=3 clones. D. Schematic for differentiation of iPSC-HSPCs to lymphoid progenitor cells and DP T cells.

**Figure 2.**
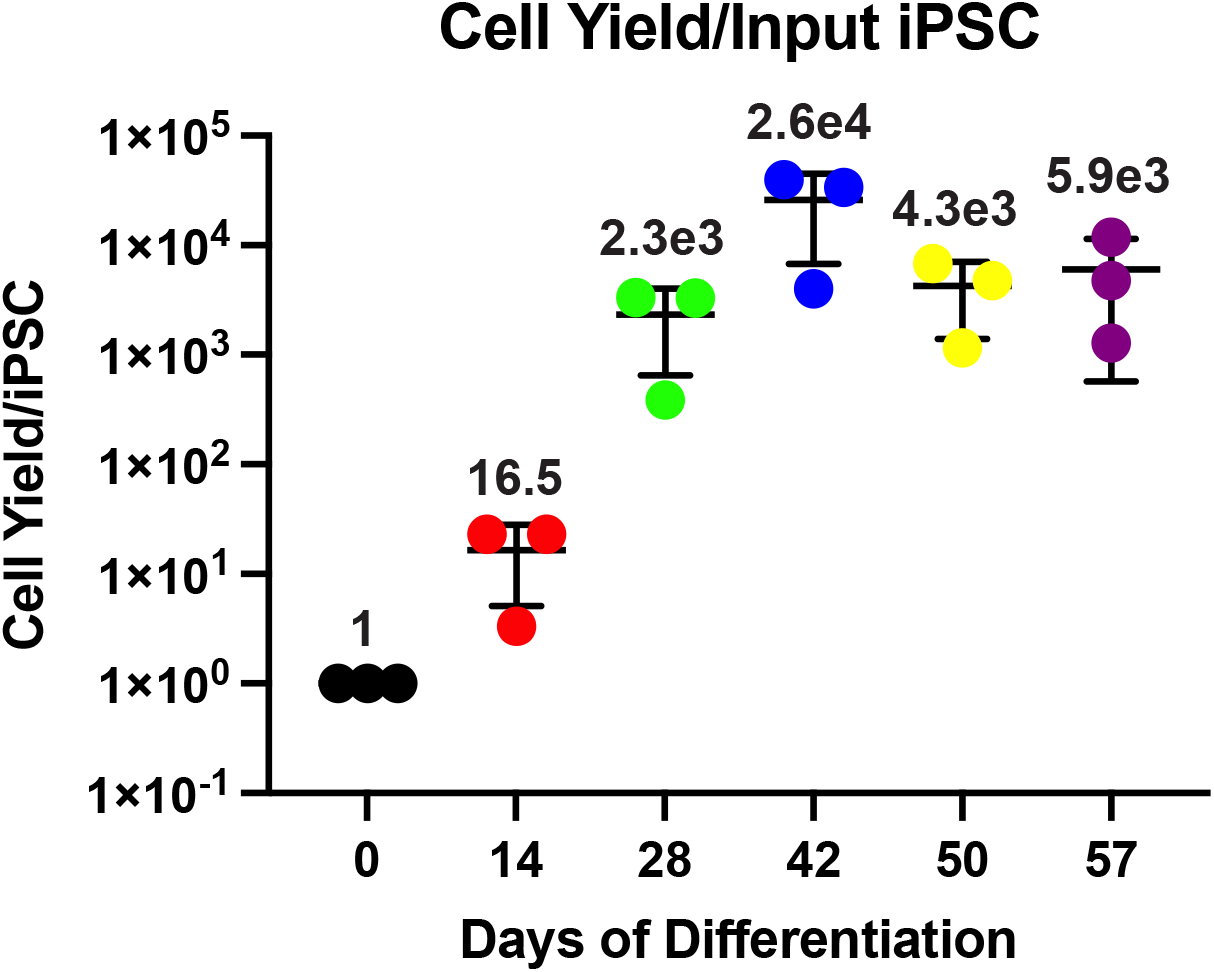
Yield of iPSC-derived hematopoietic stem and progenitor cells (HSPCs), lymphoid progenitors, double positive (DP) T cells, CD4sp T cells, and Tregs. Cell yield per input iPSC at various stages of differentiation. Data represent mean ± SD of n=3 independent experiments from n=1 clone.

For commitment and differentiation to the T cell lineage, interaction between blood cell progenitors and a Notch ligand is required^26–28^. To differentiate the iPSC-HSPCs to lymphoid progenitors, the entire population of free-floating HSPCs was separated from the EBs, collected, and seeded onto plates coated with lymphoid differentiation coating material (LDCM), a commercially available matrix containing a Notch ligand **(Figure 1D)**. The HSPCs were then further differentiated to lymphoid progenitor cells that were CD5 and CD7 DP by day 28 (**Figure 1C**, 67.4±0.1%, n=6) and subsequently to CD4 and CD8 DP T cells by day 42 (**Figure 1C**, DP T cells, 86.3±0.1%, n=5) using a commercially available iPSC-HSPC to T cell differentiation system **(Figure 1D)**. The approximate yield of lymphoid progenitor cells and DP T cells was 2.3e3±1.4e3 per input iPSC and 2.6e4±1.6e4 per input iPSC, respectively **(Figure 2)**. We tested the use of CXCL12 (SDF1a) and a p38 inhibitor (SB203580) to enhance the number of DP T cells^22^ and found an approximately 1.8-fold increase in DP T cells **(Supp. Figure 2)**.

### Generation of CD4-sp T cells and FOXP3^+^ Tregs from iPSC-CD4^+^CD8^+^ DP T cells

During T cell development in the thymus, TCR signaling is required in DP thymocytes to induce positive selection and subsequent generation of either mature CD4^+^ or CD8^+^ T cells^29^. In addition to differentiation into T cells, the interaction between the TCR and MHC molecules in the thymus also rescues thymocytes from programmed cell death^30^. Alternatively, it has been shown that DP thymocytes can undergo positive selection and maturation into CD4^+^ T cells without engagement of the TCR if provided with a strong, persistent signal^29, 31, 32^. We tested the ability of the iPSC-derived DP T cells (iPSC-DP T) to convert to CD4sp T cells by exposure to three different doses (low, medium, high) of the potent cell stimulator PMA/Ionomycin (PMA/I) **(Figure 3A).** A change in phenotype was observed by 24 hours when the iPSC-DP T cells appeared to be primarily CD8sp initially but shifted dramatically to become CD4sp 4 days after treatment. After 6 days, more than 50% of the cells were CD4sp at the lowest concentration of PMA/I **(Figure 3B)**. Medium to high concentrations of PMA/I were also effective in converting iPSC-DP T cells to CD4sp T cells albeit at lower efficiency (**Supp. Figure 3**). The cells remained DP in the absence of PMA/I. Commitment to the CD4 lineage is typically specified by the zinc-finger transcription factor and master regulator ThPOK ^33^. At the lowest concentration of PMA/I, ThPOK could be observed in the CD3^+^TCRαβ^+^CD4^+^ population of developing T cells **(Figure 3C)**. Without PMA/I stimulation, the DP T cells did not progress towards the CD4 lineage and the minor population of CD4 ^+^ cells did not express ThPOK. The yield of CD4sp cells from PMA/I treated iPSC-DP T cells was approximately 4.3e3±2.3e3/input iPSCs **(Figure 2)**. The generated iPSC-CD4sp cells expressed CD45RO^+^, but not CD45RA, and highly expressed the co-receptor CD28. CD62L was expressed at low levels compared to primary Tconv (**Figure 3D** and **Supp. Figure 4**).

**Figure 3.**
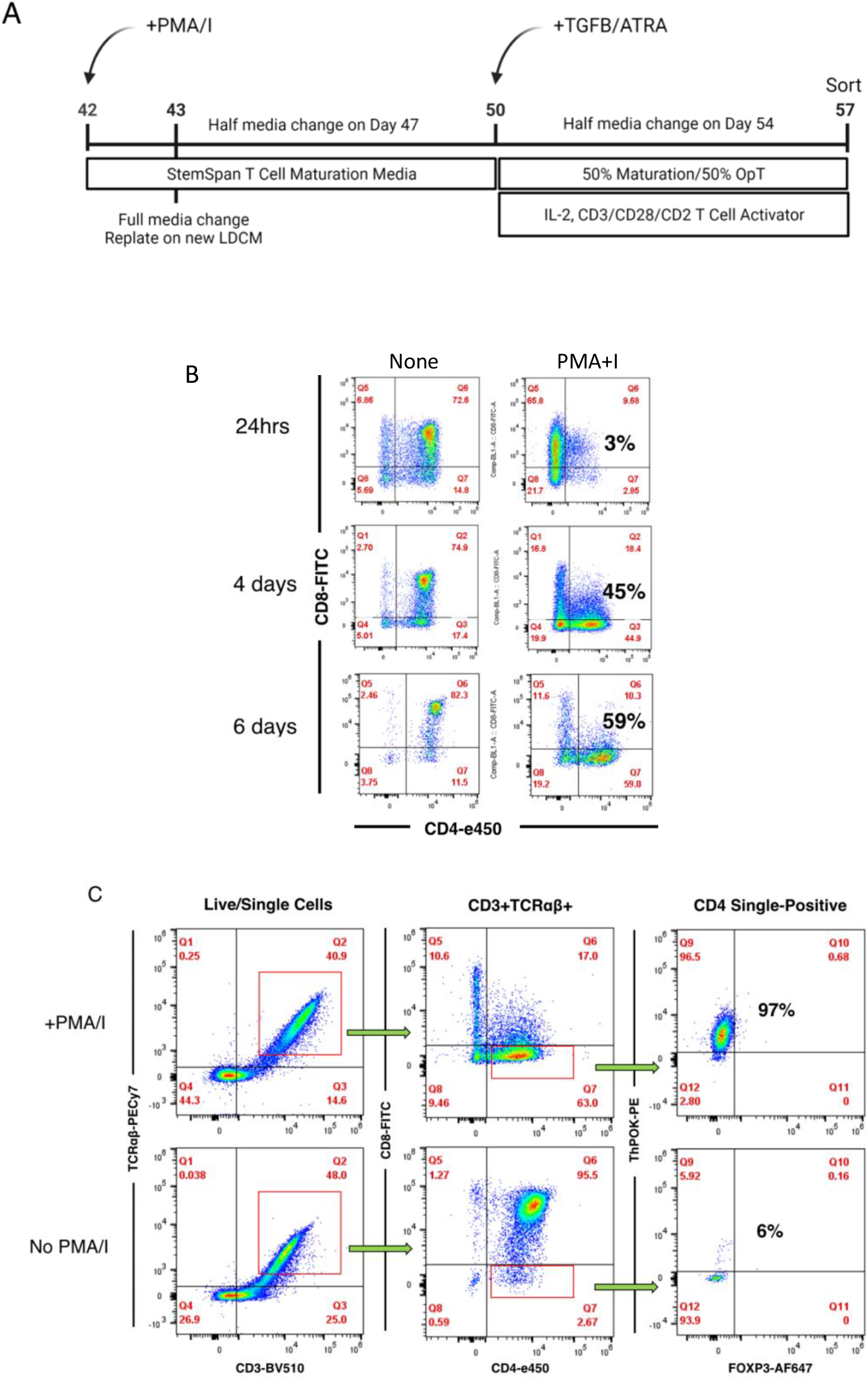
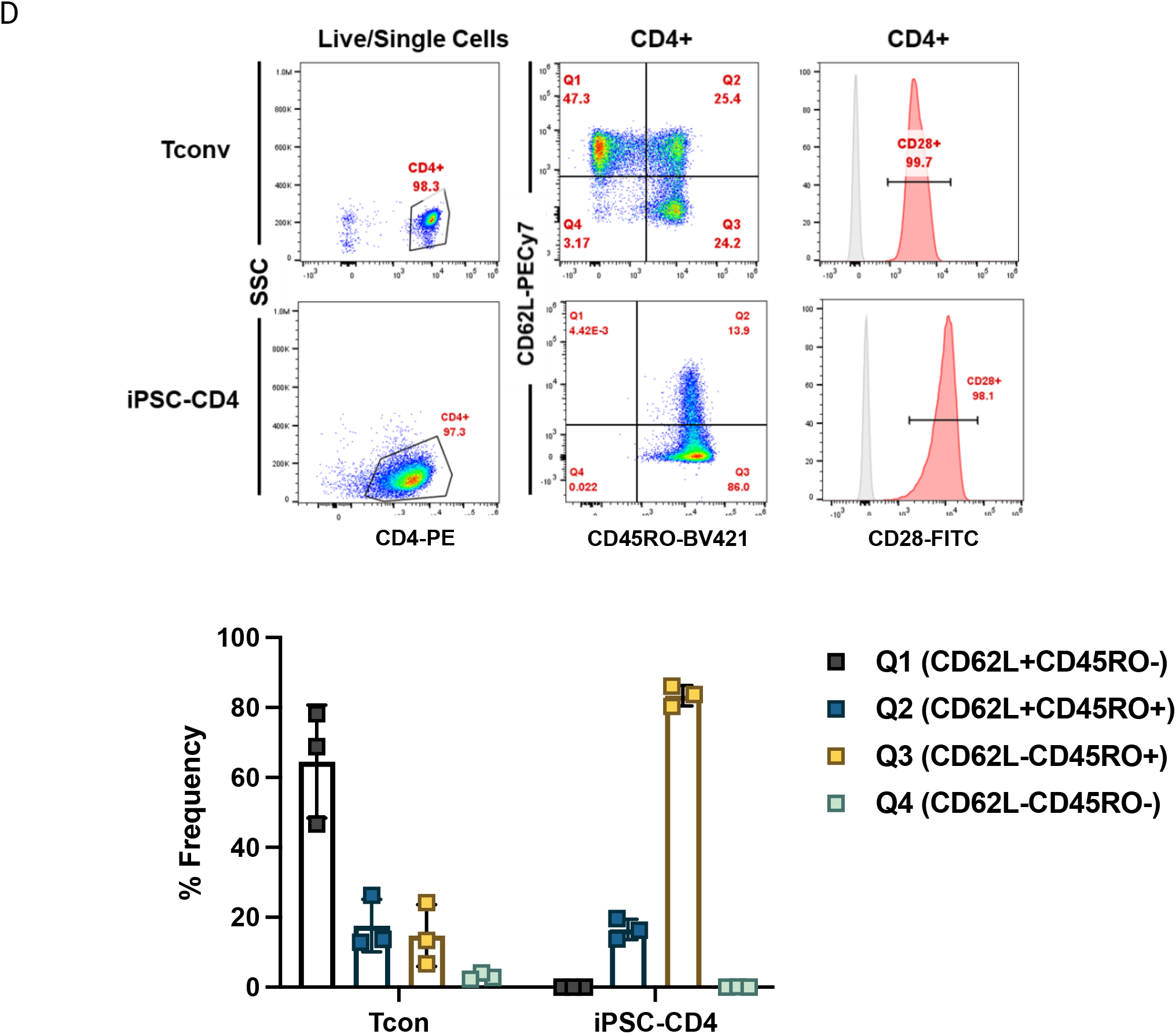
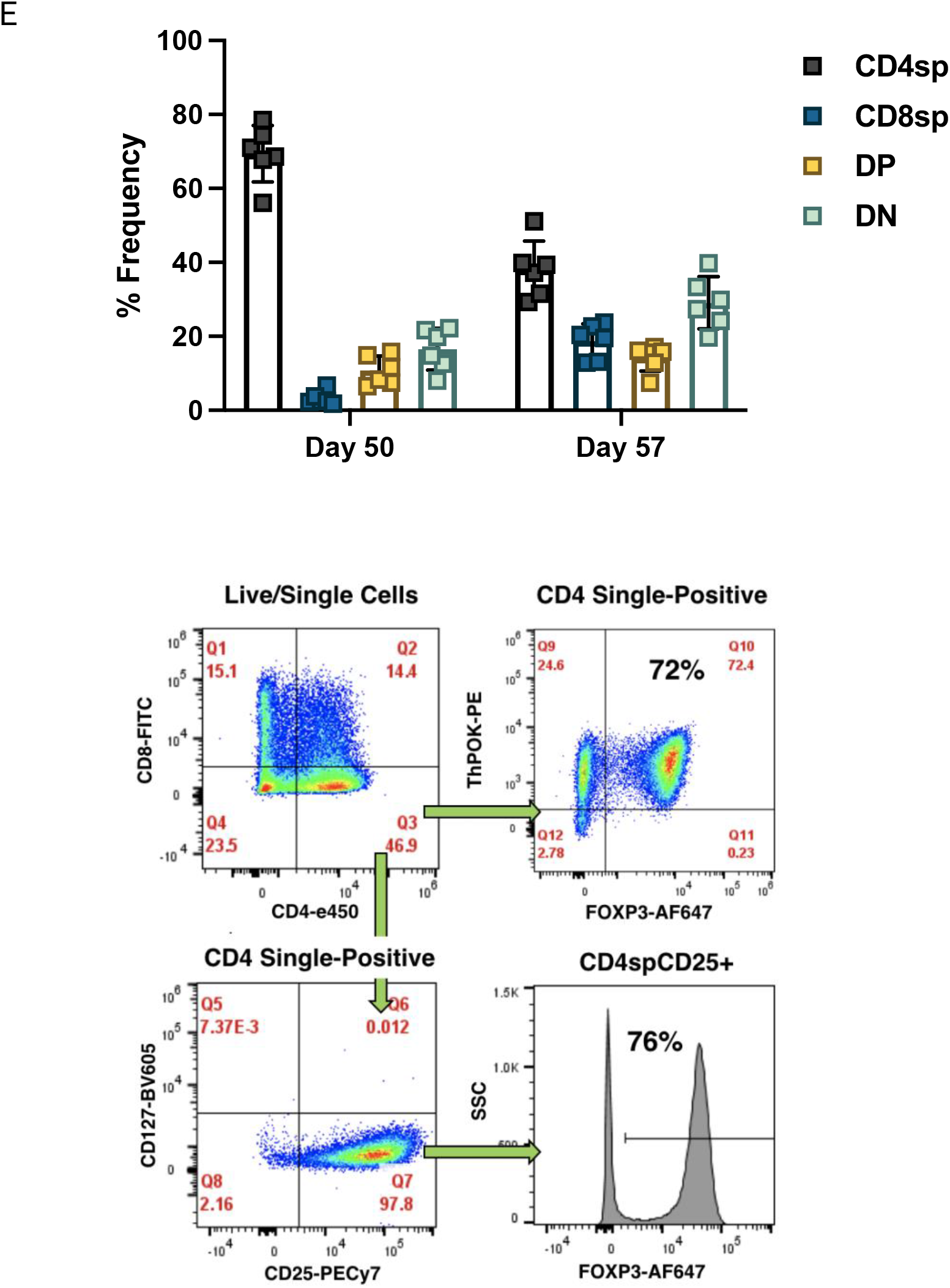
Differentiation of iPSCs to CD4sp T cells and Tregs in serum- and feeder-free conditions. A. Schematic for differentiation of iPSCs to CD4sp T cells and Tregs. B. Representative kinetic analysis of developing CD4sp T cells after Phorbol 12-myristate 13-acetate and Ionomycin (PMA/I) addition (low dose). Phenotype was assessed by the expression levels of CD4 and CD8 by flow cytometry (n=10 experiments from n=3 clones). C. Representative flow plots of iPSC-CD4sp T cells 8 days after PMA/I addition (day 50 of differentiation) to assess for CD4 commitment. Phenotype was assessed by the expression levels of CD3, TCRαβ, CD4, CD8, FOXP3 and ThPOK in cells with and without PMA/I treatment (n=6 experiments from n=3 clones). D. Representative flow plots and expression of CD4 markers in unactivated T conventional (Tconv) and sorted unactivated iPSC-CD4sp T cells. Phenotype was assessed by flow cytometric analysis (n=3 experiments from n=1 clone) in Tconv and iPSC-CD4sp at day 50 of differentiation for the expression of CD4, CD62L, CD45RO and CD28. CD28 FMO control (gray) and CD28 expression (red). E. Frequency of cell subpopulations from all live cells at day 50 and day 57 of differentiation (top) and representative flow cytometric plots of iPSC-Tregs 7 days after TGFβ+ATRA addition (day 57 of differentiation) (bottom). CD4sp, CD8sp, double positive (DP), and double negative (DN) populations were assessed by flow cytometric analysis of CD4 and CD8 expression (n=6 experiments from n=3 clones). Data show either a representative flow plot set of n independent experiments or represent mean ± SD of n independent experiments.

After successfully generating CD4sp T cells from iPSCs, we next interrogated whether a regulatory phenotype could be induced to produce Tregs. Previous studies have shown the ability to generate FOXP3^+^ Tregs from CD4^+^CD25^-^ T cells, termed induced Tregs (iTregs), in the presence of TGFβ and IL-2^34, 35^ ^36^. To determine if the iPSC-CD4sp T cells could also be induced to express FOXP3, cells were treated with a commercially available cocktail of TGFβ, all-trans retinoic acid (ATRA), and IL-2 and supplemented with a soluble T cell activator composed of anti-human CD3, CD28, and CD2 monospecific antibody complexes on day 50 of differentiation **(Figure 3A)**. Seven days post-treatment (day 57) with the TGFβ cocktail and soluble activator, an increase in frequency of DN and CD8sp T cells was observed and, while reduced, the culture retained a sizeable CD4sp T cell population **(Figure 3E)**. When gating on the CD4sp T cell population the cells expressed CD25, low levels of CD127 and high levels of FOXP3 similar to primary Tregs. Approximately 26% (+/-9.2%; n=6) of the total live population displayed a ≥90% CD4spCD25CD127lowFOXP3^+^ phenotype. In addition, the CD4sp T cell population maintained expression of ThPOK while co-expressing FOXP3 **(Figure 3E)**. We observed an average yield of 5.9e3±4.4e3 (n=3) iPSC-Treg/input iPSC **(Figure 2)**.

### iPSC-CD4sp T cells displayed a molecular signature similar to that of unconventional T cells while iPSC-Tregs displayed a molecular signature similar to that of primary Tregs

As the populations of iPSC-CD4sp cells and iPSC-Tregs are heterogenous post-PMA/I and post-TGFβ and ATRA treatment, we sought to isolate the CD4sp population at day 50 and the target Treg population of CD4spCD25^+^CD127^-^FOXP3^+^ iPSC-Tregs at day 57 by FACS to further characterize them by single-cell RNA sequencing (scRNA-seq). For iPSC-Tregs, a population of target cells that is >90% FOXP3 was obtained by gating on a “CD25 high” population from the total population of live, single CD4sp T cells **(Supp. Figure 5)**.

To determine the similarity between the iPSC-CD4sp to primary CD4 T cells and the iPSC-Tregs to primary Tregs, their transcriptomes were compared. Five cell samples were analyzed: (1) sorted iPSC-Tregs; (2) primary Tregs; (3) induced Tregs (generated from primary naïve CD4 T cells); (4) iPSC-CD4sp; and (5) primary CD4 Tconvs. A global comparison of differentially expressed genes within the 5 samples by hierarchical clustering revealed that iPSC-CD4sp T cells were similar in their transcriptional profile to primary Tconv while primary Tregs were similar to induced Tregs. The iPSC-Tregs however, were distinct in their gene expression profile **(Figure 4A).** Using Cell Typist and the Human Cell Atlas public data^13, 37, 38^ to predict and automatically annotate scRNAseq data, we show that the cells in all five samples were classified as T cells with high probability (**Supp. Figure 6A**). In addition, a large majority of primary Tregs & iPSC-Tregs were declared as Tregs by CellTypist when compared to either adult or fetal cell types (**Supp. Figure 6B** and 6C). The iPSC-CD4sp were more similar to type 3 innate lymphoid (ILC3) cells when compared to adult immune cells (**Supp. Figure 6B**) and were not able to be classified when compared to fetal immune cells (**Supp. Figure 6C**). The lower transcripts in the iPSC-CD4sp population compared to primary CD4 T cells may (**Supp. Figure 6D**) explain the difficulty in classifying these cells using Cell Typist. An analyses of marker genes across developmental stages and cell types shows that both iPSC-Tregs and iPSC-CD4sp display a more immature phenotype as compared to their primary adult counterparts **(Figure 4B)**. In addition, iPSC-CD4sp express genes that are found in innate immune cells and in unconventional fetal T cells, in particular a marked expression of *ZBTB16* (PLZF) found in fetal unconventional T cells and innate cells **(Figure 4B** and **Supp. Figure 6D)**.

**Figure 4.**
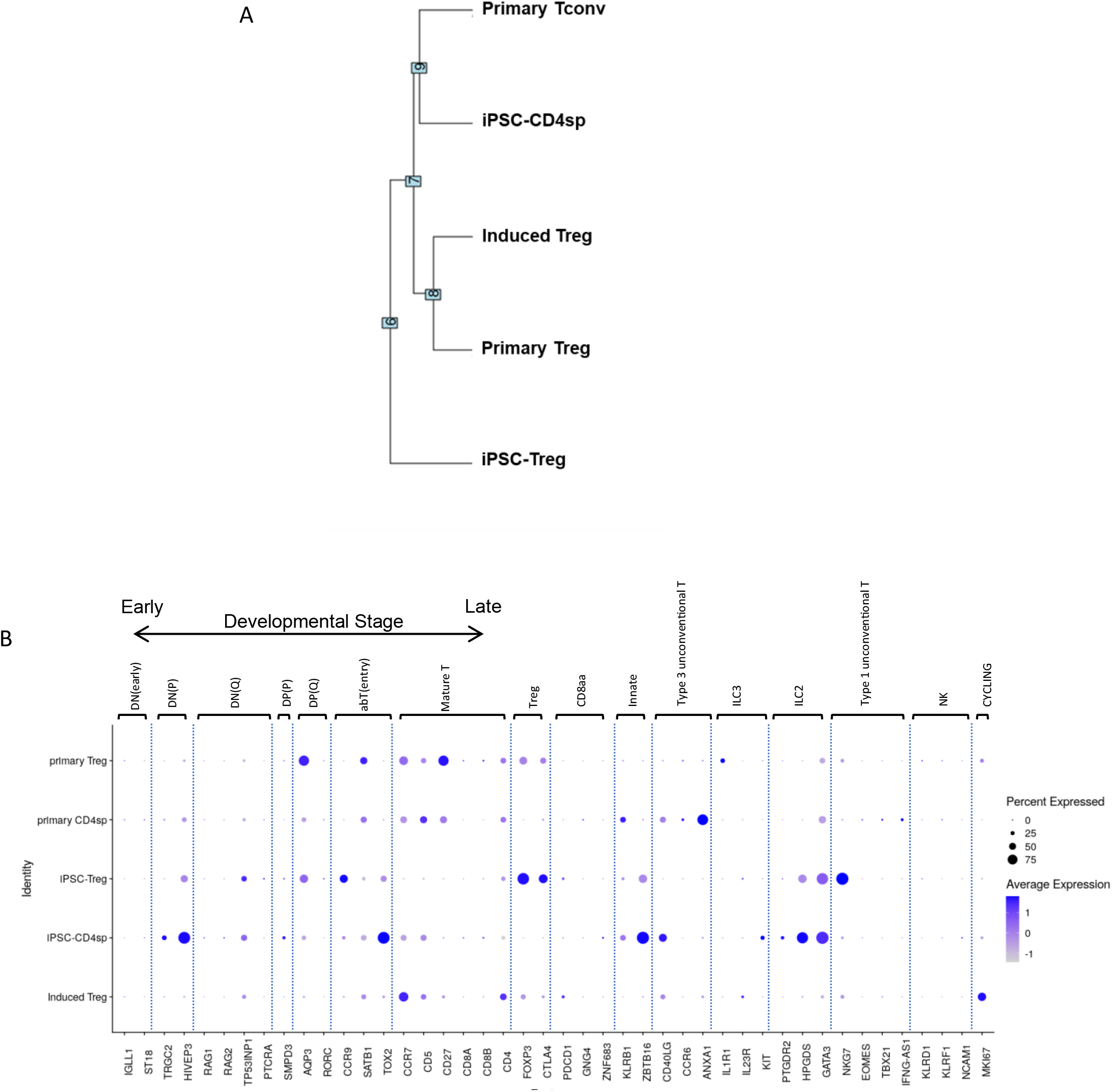
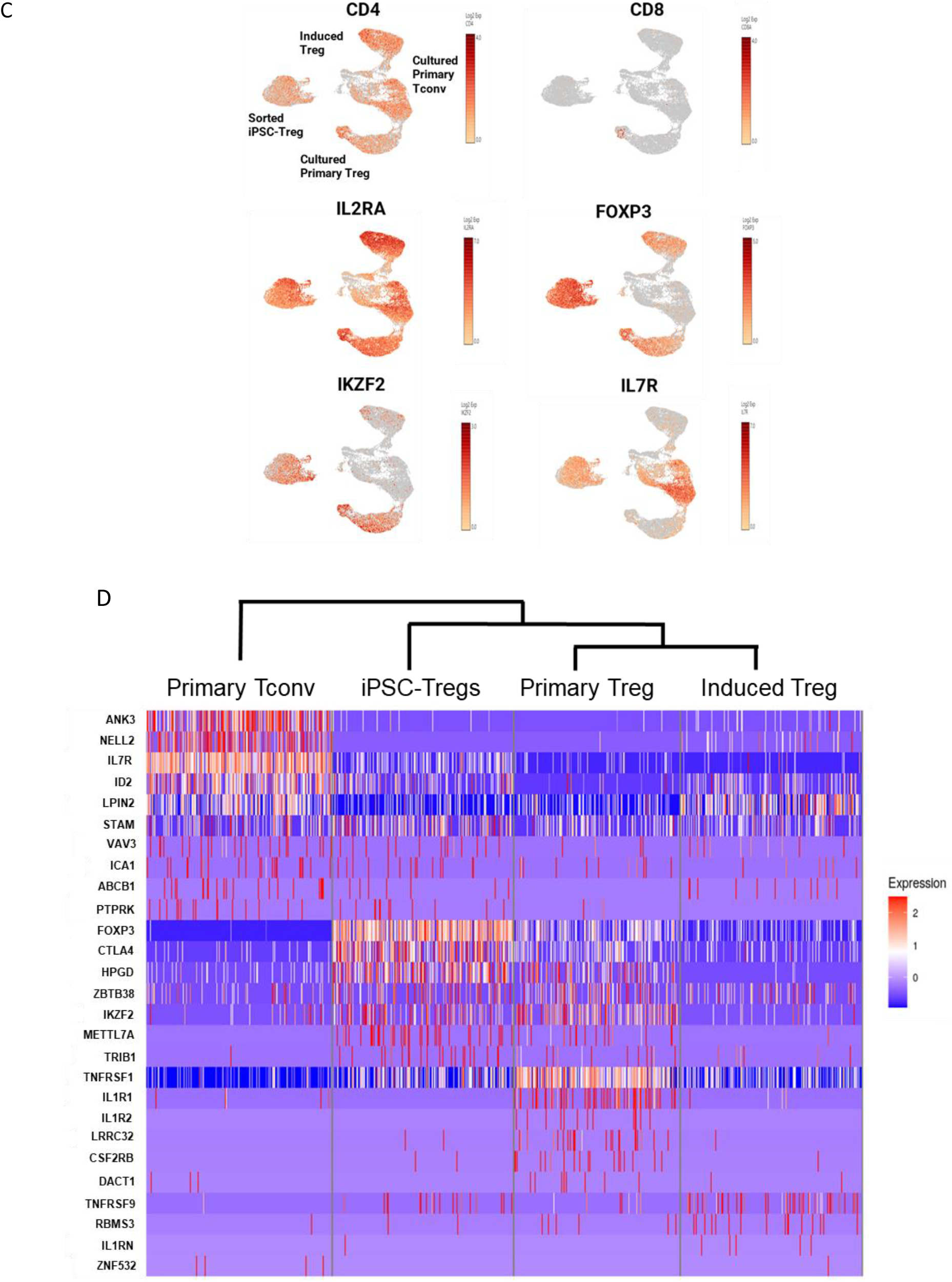
Single cell gene expression analysis of iPSC-CD4sp and iPSC-Tregs. A. Hierarchical clustering of iPSC-Tregs, iPSC-CD4sp Tcells, cultured primary T conventional (Tconv), cultured primary Treg and induced Tregs based on differentially expressed genes (DEGs). DEGs with fold change <2 and p value<0.05, were subjected to CLC analysis to perform clustering. n=1 sample for each cell type was included in the analysis. B. Dot plot of marker genes for induced Treg, iPSC-Treg, primary CD4sp, primary Treg and primary CD4sp aligned to adult immune cells. For each gene, the average normalized expression (dot color) together with the percentage of cells expressing the gene (dot size) are shown for each cluster. Annotation based on Suo et al., 2022 ^13^. *TRDC* was not found in the dataset as the iPSC cells were derived from TCRab CD4+ cells and cloned. C. Visualization of cells by cluster for iPSC-Tregs, cultured primary T conventional (Tconv), cultured primary Treg and induced Tregs using UMAPs. D. scRNA-seq heatmap showing hierarchical clustering of a core Treg gene signature in iPSC-Tregs, iPSC-CD4sp, cultured primary Tregs, induced Tregs, and cultured primary Tconv.

A visualization of cells by cluster for the iPSC-Tregs, cultured primary Tconv (CD4sp), cultured primary Treg and induced Tregs was performed using UMAPs (**Figure 4C**). Known markers of Tregs and their segregation among the clusters was explored. All clusters express *CD4* but express low to no levels of *CD8*. All clusters express *IL2RA* (CD25) with particularly strong expression in induced Tregs. *FOXP3* expression was strongest in sorted iPSC-Tregs followed by primary cultured Tregs and induced Tregs in contrast to Tconvs which do not express high levels of *FOXP3*. Both sorted iPSC-Tregs and primary Tregs express higher levels of *IKZF2* (Helios) than primary Tconv and induced Tregs. *IL7R* (CD127) is highly expressed in primary Tconv with mid- to low-level expression in all other clusters. We sought a more refined analysis and examined the expression of a core Treg gene signature in all samples^39, 40^ ^41^ **(Figure 4D)**. We found that the iPSC-Tregs were similar to primary Tregs and induced Tregs with some differences in gene expression. The iPSC-Tregs had upregulated expression of *FOXP3 and CTLA4* and downregulated expression of *TNFRSF1B*.

### Characterization and functional assessment of iPSC-CD4sp T cells and iPSC-Tregs

Further characterization and functional assessment were performed on sorted iPSC-CD4sp T cells and compared to their primary Tconv counterpart. iPSC-CD4sp T cells were activated with CD3/CD28/CD2 soluble activator. After activation, cells increased their expression of activation markers CD25 and PD-1 at 24 hours **(Supp. Figure 7A and Supp. Figure 7B)** and at 48 hours **(Figure 5A and Figure 5B)** while decreasing expression of CD28 similar to Tconv **(Supp. Figure 7B and Figure 5B)**. While CD69 MFI increased after activation, the frequency of cells expressing CD69 did not change. Activated iPSC-CD4sp T cells also secreted pro-inflammatory cytokines IL-2 and TNF-α similar to activated primary Tconv but did not secrete IFN-γ. Similar levels of IL-4 and IL-10 were secreted by iPSC-CD4sp T cells compared to primary Tconv after stimulation **(Figure 5C).**

**Figure 5.**
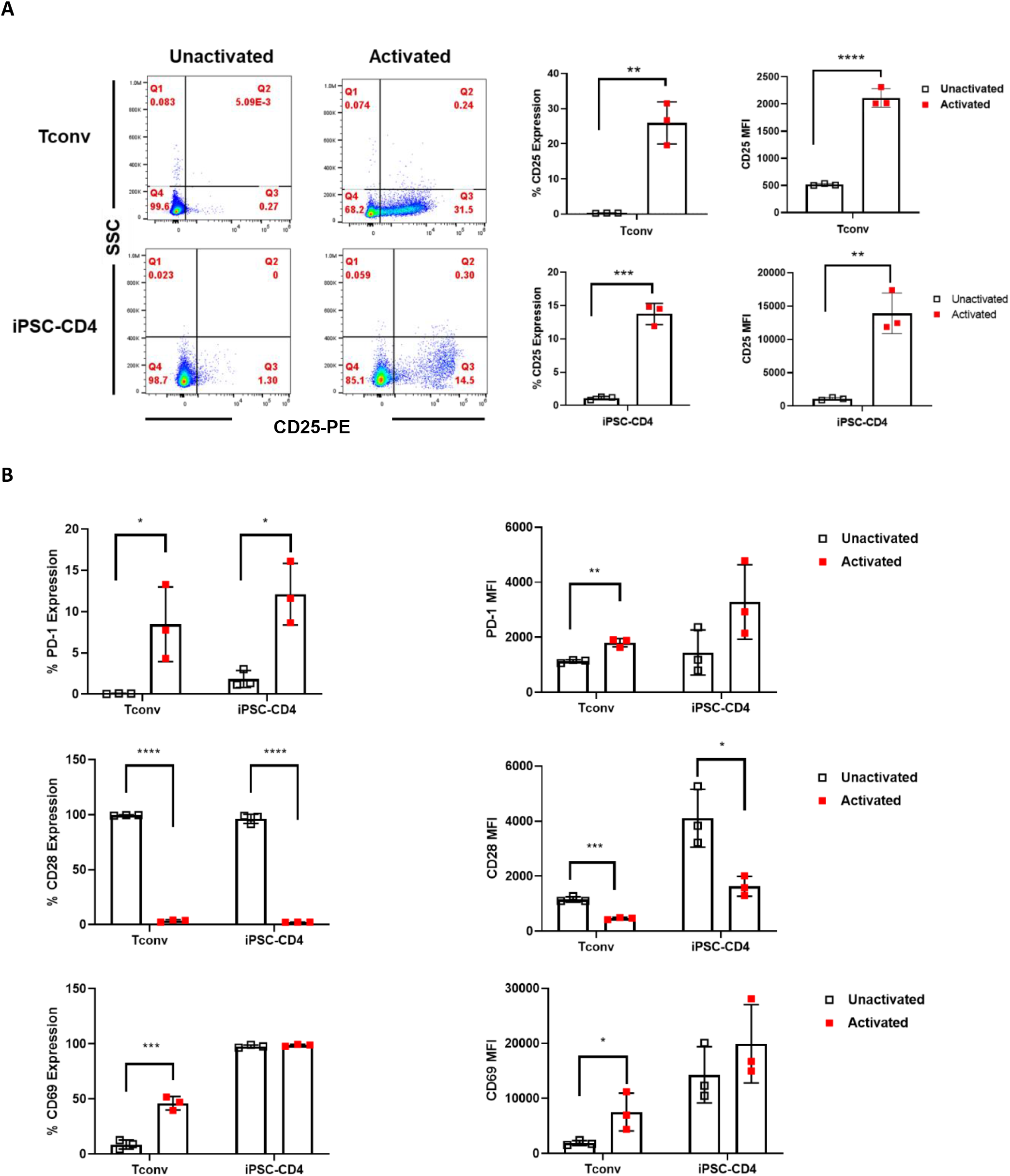
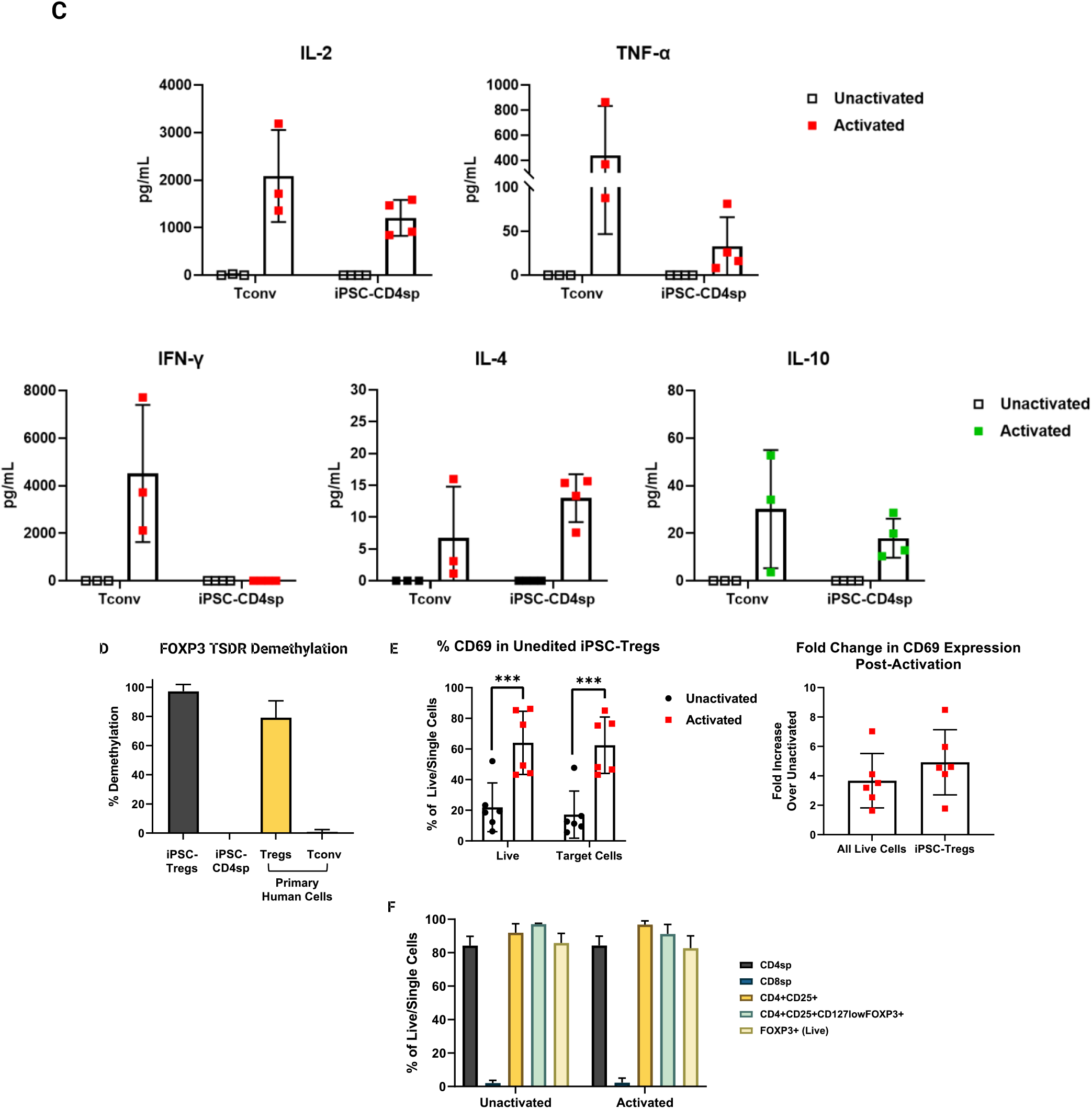
Characterization of sorted and activated iPSC-CD4sp and iPSC-Tregs. A. Representative flow plots and expression of CD25 in sorted unactivated and activated iPSC-CD4sp T cells (n=1 clone) or T conventional (Tconv) cells (n=3) by flow cytometry after 48 hours. CD25 Median fluorescence intensity (MFI) was calculated (right). B. Expression of activation markers (CD69, PD-1 and CD28) was assessed by flow cytometry after 48 hours of activation. MFI was calculated (right). C. Secretion of cytokines from sorted unactivated and activated iPSC-CD4sp T cells 24 hours after activation (n=3, iPSC-CD4sp from n=1 clone; n=3, primary Tconv). D. Assessment of FOXP3 TSDR demethylation in iPSC-Tregs, iPSC-CD4sp T cells, primary Tregs, and primary Tconv. Post-sort iPSC-Tregs were collected at day 57 for analysis and compared to primary controls (n=4, iPSC-Tregs; n=3, iPSC-CD4sp T cells, n=3, primary Tregs; n=3, primary Tconv). E. Expression of CD69 in sorted unactivated and activated iPSC-Tregs. The expression of CD69 was assessed in both total live cell populations and target iPSC-Tregs by flow cytometry (left). Fold change in CD69 expression was assessed in activated total live and target iPSC-Tregs compared to unactivated total live cells and iPSC-Tregs (right, n=6 from n=3 clones, unactivated and activated). F. Phenotype of unactivated and activated iPSC-Tregs. Treg markers were assessed by flow cytometric analysis (n=6 from n=3 clones). Data represent mean ± SD of n independent experiments. P*<0.05, P**<0.01, P***<0.001, P****<0.0001 by unpaired two-tailed t-test.

We then characterized and examined the *in vitro* function of sorted iPSC-Tregs. Immunophenotyping of cells post-sort showed a typical Treg phenotype with an average of 83.4±6.3% FOXP3 expression (n=5) and average 91.6±1.6% Helios expression (n=2) from total live cells. Demethylation of the evolutionarily conserved non-coding element FOXP3 Treg-specific demethylated region (FOXP3 TSDR) is required for stable FOXP3 expression in Tregs^42^. It is typically methylated in Tconv, induced Tregs and other non-Treg blood cells^43^. We examined the methylation state of FOXP3 TSDR by high resolution melt quantitative polymerase chain reaction (HRM-qPCR) in the sorted iPSC-Tregs. The iPSC-Tregs demonstrated >90% demethylation similar to the primary Treg positive control. In contrast, primary CD4^+^CD25^-^ Tconv as well iPSC-CD4sp T cells were highly methylated at FOXP3 TSDR **(Figure 5D)**. The sorted iPSC-Tregs could be activated via their TCR using beads coupled to anti-CD3 and anti-CD28 antibodies (Dynabeads). After activation, the iPSC-Tregs increased the expression of early activation marker CD69 on average of 3-fold with no change to their Treg phenotype **(Figure 5E-F)**.

Tregs typically exert their suppressive function through various mechanisms including suppression by cytolysis^44, 45^. Cellular toxicity was determined by co-culturing activated and unactivated iPSC-Tregs with primary CD4 Tconv target cells. Compared to activated cytotoxic primary CD8 T cells, activated iPSC-Tregs had low cytolytic activity similar to activated primary Tregs as measured by LDH release assay **(Figure 6A).**

**Figure 6.**
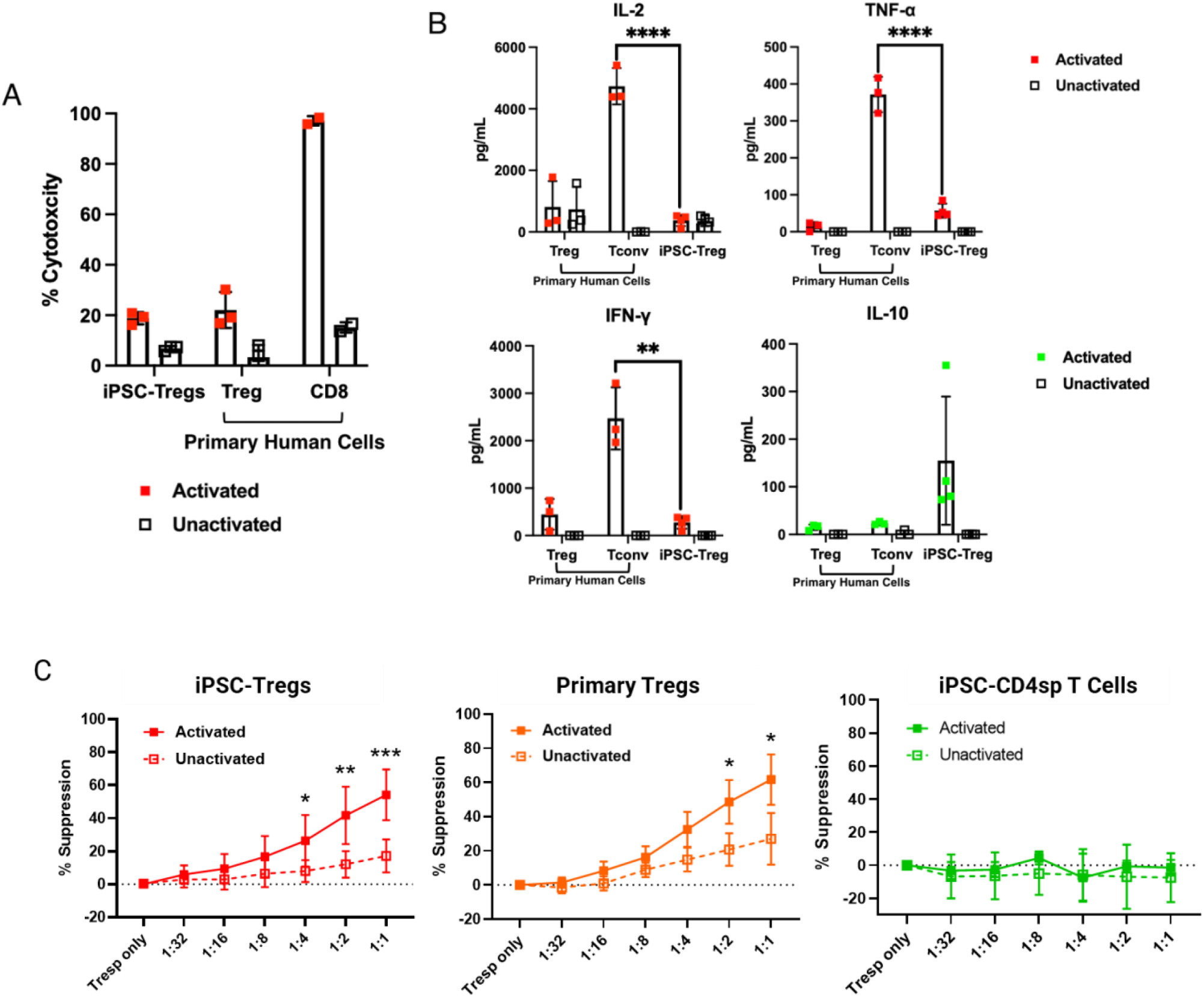
Functional analysis of iPSC-Tregs. A. Cytotoxic activity of sorted unactivated and activated iPSC-Tregs as measured by an LDH release assay. iPSC-Tregs, primary Tregs, and primary CD8 T cells were cocultured with T conventional (Tconv) cells and LDH was measured after 3 days (n=3 from n=1 clone, iPSC-Tregs; n=3, primary Tregs; n=2, primary CD8 T cells). B. Secretion of cytokines from sorted unactivated and activated iPSC-Tregs 24 hours after activation (n=4 from n=1 clone, iPSC-Tregs; n=3, primary Tregs; n=3, primary Tconv). C. Suppression of Tconv proliferation by unactivated and activated iPSC-Tregs, iPSC-CD4sp T cells and primary Tregs. iPSC-Tregs, primary Tregs, and iPSC-CD4sp T cells were cocultured with activated CTV labeled Tconv and CTV dilution was measured by flow cytometry after 3 days (n=6 from n=3 clones, iPSC-Tregs; n=4, primary Tregs; n=3 from n=3 clones, iPSC-CD4sp). Data represent mean ± SD of n independent experiments. P*<0.05, P**<0.01, P***<0.001 by unpaired two-tailed t-test.

Additional suppressive mechanisms include secretion of anti-inflammatory cytokines like IL-10^44^. IL-10 is produced by a variety of cells including CD4^+^ Th2 cells, Tregs and IL-10-induced CD4^+^ T reg cells (Tr1 cells)^46^. Interestingly, iPSC-Tregs express higher, but variable, amounts of IL-10 as compared to primary Tregs. Importantly, the iPSC-Tregs secreted low levels of pro-inflammatory cytokines IL-2, IFN-γ, and TNF-α before and after activation, similar to primary Tregs **(Figure 6B)**.

The cardinal feature of Tregs is their ability to suppress the activation and proliferation of Tconv or Teff cells by a cell-to-cell contact mechanism^44^. To determine the suppressive function of activated iPSC-Tregs, the cells were co-cultured with activated primary Tconv and their proliferation was measured by flow cytometric analysis. Activated iPSC-Tregs effectively suppressed Tconv proliferation at a 1:1 ratio of Tregs:Tconv similar to activated primary Tregs **(Figure 6C)**. Without activation, iPSC-Tregs had significantly lower levels of suppression at 1:1 to 1:4 Tregs: Tconv ratios. An iPSC-Treg dose-dependent effect on Tconv suppression was also observed similar to primary Tregs. As expected, when iPSC-CD4sp T cells were tested for suppressive function, they were unable to prevent Tconv proliferation even at the highest ratio of 1:1 CD4sp T cells:Tconv **(Figure 6C)**. This confirms that the suppressive function is specific to iPSC-Tregs and appears only after conversion of iPSC-CD4sp T cells to FOXP3^+^ Tregs and after cell activation.

### Generation and functional assessment of HLA-A2 CAR^+^ iPSC-Tregs

We next investigated the function of iPSC-Tregs equipped with a chimeric antigen receptor (CAR). To this end, we genetically engineered iPSCs to express an HLA-A2 CAR using zinc-finger nucleases (ZFNs) targeting exon 2 of the *TRAC* gene.

We sought to retain expression of the *TRAC* gene and utilized a donor sequence comprised of *TRAC* cDNA encompassing the *TRAC* exonic sequence downstream of the integration site. This donor sequence is placed upstream and in-frame with a T2A coding sequence such that the engineered locus retains expression of the TCRα chain while expressing an HLA-A2 CAR under the control of the *TRAC* promoter **(Figure 7A)**. Through optimized electroporation of the iPSCs with TRAC ZFN mRNA and donor DNA we achieved a >80% targeted integration without selection as assessed by sequencing of the bulk edited population.

**Figure 7.**
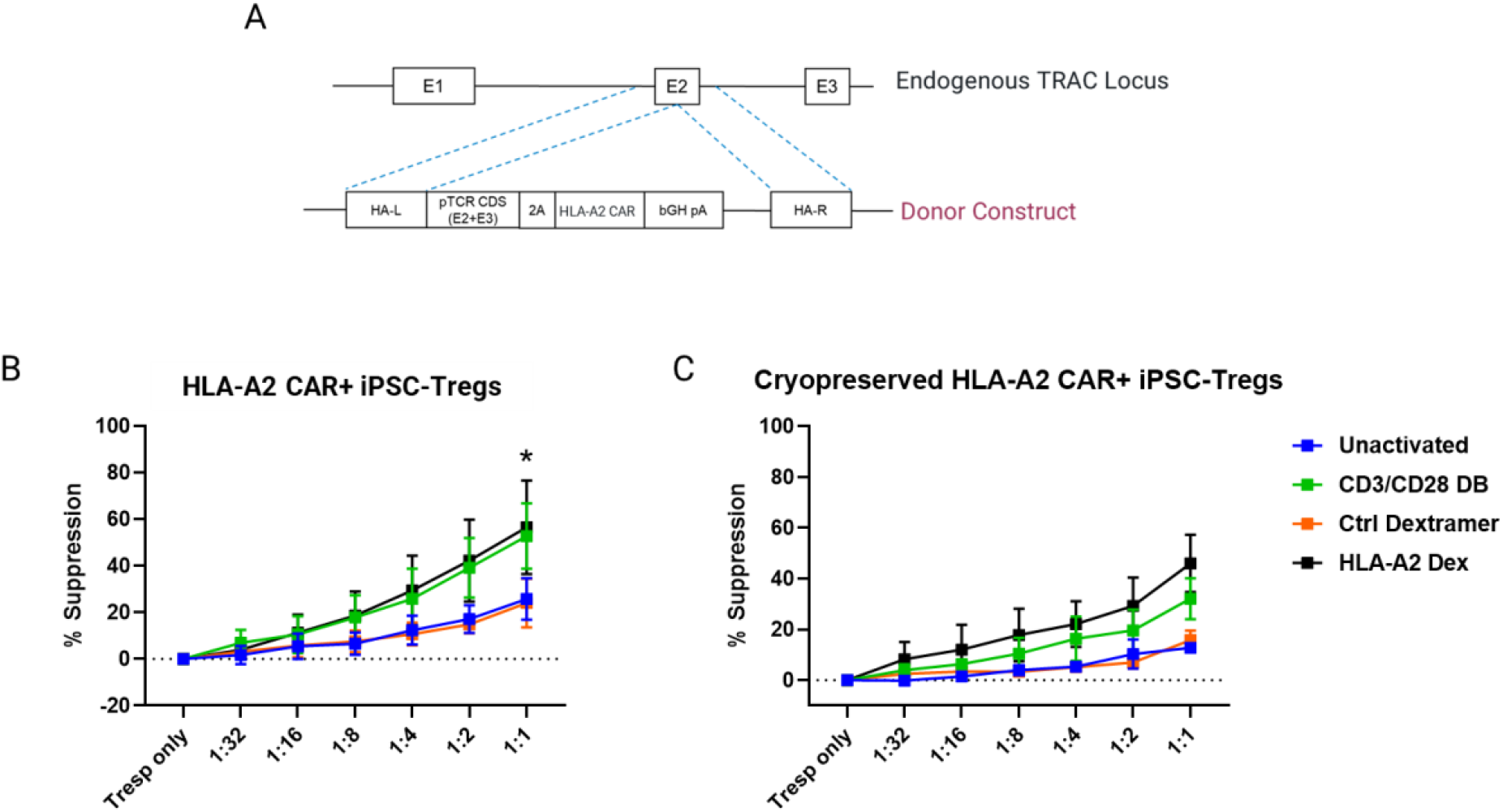
Generation and characterization of HLA-A2 CAR iPSC-Tregs. A. Editing strategy for generation of HLA-A2 CAR iPSCs. A donor construct showing the TCR coding sequence and HLA-A2 CAR sequence flanked by homology arms targets exon 2 of the endogenous T Cell Receptor Alpha Constant (TRAC) locus. B. Suppression of T conventional (Tconv) cell proliferation by HLA-A2 CAR+ iPSC-Tregs activated *via* the TCR or CAR. HLA-A2 CAR+ iPSC-Tregs were cocultured with activated CTV labeled Tconv and CTV dilution was measured by flow cytometry after 3 days (n=4 from n=2 clones). *p<0.05 C. Suppression of Tconv proliferation by cryopreserved HLA-A2 CAR+ iPSC-Tregs activated via the TCR or CAR. HLA-A2 CAR+ iPSC-Tregs were cocultured with activated CTV labeled Tconv and CTV dilution was measured by flow cytometry after 3 days (n=2 from n=2 clones). Data represent mean ± SD of n independent experiments. P*<0.05 by unpaired two-tailed t-test.

The resulting engineered iPSCs were cloned, expanded, and sequenced to obtain a biallelically modified iPSC clone with the HLA-A2 CAR inserted at the target site. Sequencing analysis confirmed the precise and in-frame insertion of a single copy of donor DNA (HLA-A2-CAR), without any detected mutations. The CAR-engineered iPSCs also displayed a normal pluripotent phenotype and were karyotypically normal **(Supp. Figure 1)**. To generate HLA-A2 CAR^+^ iPSC-Tregs (CAR iPSC-Tregs) the iPSCs were differentiated to HSPCs, lymphoid progenitor cells, and DP T cells using the same differentiation process utilized for unedited iPSCs as described above **(Supp. Figure 8A)**. HLA-A2 CAR iPSCs could also generate CD4sp T cells and FOXP3^+^ Tregs **(Supp. Figure 8B, 8C)**. Of note, we observed a 2.2-fold lower yield of Tregs (n=3, ±0.2 fold) from the HLA-A2 CAR iPSCs compared to Tregs generated from unedited iPSCs. By flow cytometric analysis, we found that ∼90% (90.1%±1.8%, n=4) of the DP T cells were already expressing the CAR by day 42 and retained high CAR expression at both the CD4sp T cell (99%±0.2%, n=4) and Treg stages (99.85%±0.1%, n=4). The intensity of CAR expression increased as the cells differentiated as measured by MFI **(Supp. Figure 8D)**.

Similar to unedited iPSC-Tregs, HLA-A2 CAR^+^ iPSC-Tregs could also be sorted. Four days post-sort the expression levels of the CAR remained stable (81.3±12.4% of TCRαβ^+^ cells and MFI=9814.3±1143.3, n=3) and the cells maintained a high level of demethylation at FOXP3 TSDR (88%±10.3%; n=3).

Further characterization showed that the CAR iPSC-Tregs could be activated through both the TCR and the HLA-A2 CAR demonstrating the engineering strategy successfully retained TRAC expression while integrating a functional CAR. Activation of the CAR iPSC-Tregs using either CD3/CD28 beads (Dynabeads) or HLA-A2 dextramer (as previously described^6^) upregulated CD69 expression compared to unactivated or dextramer control activation **(Supp. Figure 9A, 9B)**. TCR- and CAR-mediated activation induced a very low level of pro-inflammatory cytokines while inducing higher levels of IL-10 **(Supp. Figure 9C)** similar to levels of cytokines produced by unedited iPSC-Treg upon TCR activation **(Figure 6B).** The CAR iPSC-Tregs significantly suppress Tconv proliferation at similar levels to wild-type iPSC-Tregs when activated either through the TCR or the HLA-A2 CAR, while unactivated or control dextramer activated CAR iPSC-Tregs only displayed a basal level of suppression **(Figure 7B).**

### Characterization and functional assessment of cryopreserved CAR iPSC-Tregs

A requirement for a viable CAR iPSC-Treg therapy is the ability to efficiently cryopreserve and store the cells for future use without loss of stability and function. Towards that goal, we cryopreserved iPSC-Tregs post-sort and then thawed, characterized, and tested their suppressive function. Approximately 80% of cells were recovered post-thaw (avg=82.5%, n=2) with a viability comparable to pre-cryopreservation (avg=74%, n=2). The CAR iPSC-Tregs maintained their phenotype post-thaw **(Supp. Figure 10A)**. The cells were also highly demethylated at FOXP3 TSDR (95.5±0.5%, n=2) and displayed an average 4-fold increase in CD69 after activation *via* the HLA-A2 CAR **(Supp. Figure 10B).** Finally, thawed CAR iPSC-Tregs were able to suppress Tconv proliferation when activated *via* the TCR or CAR **(Figure 7C).**

## Discussion

We report here for the first time a serum- and feeder-free method to produce CD4sp T cells and functional Tregs with and without a CAR from iPSCs. While the generation of CD4sp T cells from iPSCs has been reported^10, 14^, it currently requires the use of murine stromal cells and animal serum (fetal bovine serum, FBS) in ATOs. The use of xenogenic feeders, animal serum and 3D organoids is not ideal for the development of a clinical-grade therapeutic product. This is due to issues with safety, scalability and serum batch-to-batch heterogeneity as well as difficulty dissociating cells from 3D organoids. In addition, automation of 3D organoid generation to produce large-scale reproducible organoids is a nascent field and will require further optimization to minimize variability across differentiation batches^47^. The method proposed here circumvents many of these issues and increases its amenability for manufacturing cell therapies at scale.

The reprogramming of T cells to pluripotency and subsequent differentiation to mature T cells has been well documented particularly towards the generation of CD8αβ^+^ CTLs^48, 49^ ^50^. Using T cell-derived iPSCs may aid in more efficiently generating iPSC-T cells as the early stages of T cell development that are characterized by failed TCR rearrangement and cell death are bypassed^51^. While antigen specificity is unknown, reprogramming from a T cell clone possessing a single TCR^9, 21, 52^ or introducing a CAR or transgenic TCR at the iPSC or T cell stage^8, 11, 12, 22, 53, 54^ can help confer specificity. Based on these findings, we sought to utilize a similar strategy to develop CD4sp T cells and Tregs from iPSCs. We were able to successfully obtain iPSCs from mature CD4sp T cells that expressed classic pluripotency markers and appeared as compact colonies with well-defined borders. The iPSCs were amenable to genome engineering for the targeted integration of an HLA-A2 CAR. Both the wild-type and engineered iPSC lines were karyotypically normal **(Supp. Figure 1)**. We did not perform deep characterization of off-targets in edited iPSCs at this research-grade stage. Most importantly, an advantage of using iPSCs over primary cells is the ability to select through whole-genome sequencing (WGS) well-characterized clones with no specific off-targets and expand them to generate a MCB for future clinical use.

Methods to differentiate iPSCs into HSPCs and progenitor T cells are readily available, but many require the combination of multiple individual reagents from various sources which may contribute to increased variability in differentiation. We found that utilizing commercially available reagents specifically designed to generate HSPCs and T cell progenitors from iPSCs streamlined the differentiation process while satisfying the serum- and feeder-free requirements. We utilized a modified EB-based method to differentiate iPSCs and employed commercially available serum-free media for the generation of CD34^+^CD45^+^ HSPCs capable of differentiation to lymphoid cells. It is worth noting that while the process yielded a mixture of CD34^+^ and CD34^-^ cells, no enrichment or purification for CD34^+^CD45^+^ target cells was required for further differentiation. Indeed, we found that by separation of the free-floating progenitor cells from the main EB we were able to generate lymphoid progenitor cells that co-expressed CD5 and CD7 marking their T cell commitment^55, 56^. Continued differentiation of the lymphoid progenitor cells resulted in CD4 and CD8 DP T cells again, without enrichment of the starting population.

Thymic DP T cells express both CD4 and CD8 co-receptors that promote TCR engagement of MHC ligands to enhance signal transduction^29^. Through various strengths of binding affinity to MHC ligands, the developing T cells will undergo positive selection to survive or cell death by negative selection^57^. The step-by-step natural process followed by DP T cells after positive selection to commit to either the CD4sp or CD8sp lineage is unclear^29, 58^. It has been suggested that a persistent and potentially stronger TCR signal is required to make the CD4 choice and block re-expression of the CD8 co-receptor. To mimic the signal required for positive selection and commitment to the CD4 lineage at the DP T cell stage, we utilized a strategy previously shown to induce CD4 commitment in murine DP T thymocytes without TCR engagement^31^. The addition of PMA/I activates protein kinase C and elevates calcium levels, respectively and in a similar manner to TCR-mediated activation with signals from co-stimulator molecules like CD4, CD8 and LFA-1, by-passing TCR engagement^31, 59^. At levels between 0.016 μM to 0.16 μM of PMA and 0.13 μM of ionomycin, TRAC KO murine DP T thymocytes are positively selected to CD4 T cells. Higher concentrations of PMA/I (1.6 μM and 1.3 μM, respectively) were inefficient in generating CD4 T cells. These findings are consistent with our method in that the lower concentrations we used successfully converted cells to the CD4 lineage while higher concentrations of PMA/I were less efficient. Importantly, after conversion to CD4sp T cells, ThPOK expression was observed further confirming its lineage commitment (**Figure 3C**). The kinetics of co-receptor expression in our system is different from what has been previously published in mouse-derived thymocytes^31, 59, 60^ and could be due to species-specific differences between mouse and human thymocyte development^61^, differences in the systems used (iPSC-derived vs. thymus-derivates DP T cells) or related to the transient expansion of a highly proliferative, immature intermediate (CD4^−^ CD8^+^ Mki67^+^)^61^. Further characterization of the intermediate stages will shed light on the mechanisms of positive selection induced by PMA/I in our system.

Initial characterization of iPSC-CD4sp T cells revealed interesting similarities and differences to their primary T cell counterparts. The features observed in the iPSC-CD4sp cells may be due to low levels of CD4 transcripts caused by the activated state of these cells compared to primary CD4 T cells. Indeed, the cells maintained high levels of CD69 expression prior to activation when compared to primary CD4 T cells (**Figure 5B**). Helper T cells often downregulate CD4 expression upon chronic stimulation which may explain the reduced transcripts caused by exposure to PMAI/I treatment^62^. Using CITEseq in combination with scRNAseq in the future will achieve a better characterization of the iPSC-CD4sp identity than using scRNAseq alone^63^.

The expression of *ZBTB16* (PLZF), the predominantly memory phenotype (CD45RO^+^CD62L^-^) and secretion of IL-4 observed in the iPSC-CD4sp T cells may be due to thymocyte-thymocyte (T-T) interactions that can generate CD4^+^ T cells with similar characteristics through a pathway originally described *in vitro*^64^ and subsequently evidenced in MHC class II transactivator (CIITA) transgenic mice^65^ and human fetuses^66^. A study generating iPSC-CD8 T cells through an ATO also showed high levels of *ZBTB16* and similarities to fetal type 1 innate-like T unconventional cells, which are CD8^+^, as classified by Cell Typist^13^, suggesting that *in vitro* differentiation systems such as ATOs that lack thymic epithelial cells (TECs) give rise to innate-like T cells resembling fetal unconventional T cells. While the low levels of *CD4* transcripts did not allow a clear classification of our iPSC-CD4sp T cells, their phenotype and transcriptional profile is evocative of the innate and fetal unconventional T cells generated through T-T interactions^13^. Modification of the differentiation process and integration of signals similar to those induced by TECs through culture media optimization and/or cell engineering^67^ might allow for the generation of CD4 T cells with a more mature and less innate phenotype.

We did not detect IFN-γ from the iPSC-CD4sp T cells but observed secretion of pro-inflammatory cytokines IL-2 and TNF-α upon stimulation, as well as Th2-like IL-10 and IL-4 cytokines^68^ ^15^. This profile is similar to fetal unconventional CD4^+^ PLZF^+^ T cells which, when stimulated, upregulate a variety of cytokines not usually produced by a single Th subset^69^. Different levels of TCR activation induce different cell fates and modifying the TCR activation levels might modify the cell fate towards a Th1 vs. Th2 phenotype^70^.

In addition to maintaining a serum- and feeder-free differentiation system, the use of PMA/I to generate iPSC-CD4sp T cells is an especially critical finding as providing a positive selection signal towards the CD4 lineage is the first step to generating Tregs. After maturation, the iPSC-CD4sp T cells were treated with TGFβ, ATRA, IL-2 and a soluble CD3/CD28/CD2 activator to convert them to a FOXP3^+^ Treg phenotype. The generation of Tregs from naïve CD4 T cells using TGFβ is well-documented^36^. While TGFβ-induced iTregs express FOXP3, they are only partially demethylated at FOXP3 TSDR and are unstable in their expression of FOXP3. In addition, polyclonal iTregs do not always display a regulatory phenotype, are not always suppressive and can produce high levels of inflammatory cytokines^71^. In contrast, the iPSC-Tregs generated within our system were highly demethylated at FOXP3 TSDR, suppressed Tconv proliferation and did not produce inflammatory cytokines despite utilizing TGFβ towards its conversion. Transcriptomic analysis by scRNA-seq further confirmed the identity and similarity of the iPSC Tregs to primary Tregs as assessed by Cell Typist. The iPSC-Tregs expressed typical markers of their primary natural Treg counterpart, namely highly expressing *FOXP3*, *IKZF2* and *IL2RA* in the absence of activation. The iPSC-Tregs also expressed a previously published^12^ core Treg signature^39–41^ similar to that of primary Tregs and less similar to iTregs and Tconv. Differential expression of genes between iPSC-Treg and primary Treg, such as *IL7R* and *ID2*, could be due to culture conditions or to differences between cells that differentiated *in vitro* as compared to *ex vivo* adult cells. Finally, since scRNA-seq utilizes relative expression levels, measuring the absolute expression of these markers by bulk RNA-seq may provide better quantitative results to distinguish these populations.

Previous studies have shown that iPSCs derived from T cells express TCRαβ prematurely which interferes with DP T cell formation and can be overcome by supplying the cells with a strong Notch signal^11^. Constitutive expression of CAR with simultaneous expression of the endogenous TCR, however, offsets the effects of a stronger Notch signal. In this case, both elimination of TRAC expression as well as attenuation of the CAR signal was required to restore DP T cell generation^11,12^. These are important considerations for the generation of a scalable cell therapy product using engineered iPSCs. While we were able to generate DP T cells from both wild-type and engineered iPSCs, the exact Notch ligand in the lymphocyte differentiation coating material (LDCM) reagent is unknown making the need to test a known Notch ligand necessary for improving DP T cell yields. In addition, we observed a lower yield of HLA-A2 CAR^+^ iPSCs-Tregs as compared to Tregs generated from unedited iPSCs, supporting the idea that premature TCR expression and CAR signaling influences DP T cell yields and subsequent Treg generation. Defining the Notch ligand, optimizing CAR signaling as well as applying compound-based measures to improve DP T cell yield from iPSCs can aid in the generation of a more robust and reproducible differentiation protocol ^11^.

Cryopreservation is key to the development and delivery of successful cell therapy products. We found that our CAR iPSC-Tregs were amenable to cryopreservation and showed similar viability pre- and post-cryopreservation with approximately 80% recovery post-thaw. The phenotype of the cells was also maintained pre- and post-thaw with appreciable upregulation of CD69 post-activation via the CAR or TCR. Importantly, the cryopreserved iPSC-Tregs suppress Tconv proliferation in vitro confirming that the cells maintain functionality **(Figure 7)**.

In summary, we demonstrate here a method to generate functional and stable iPSC-derived Tregs and CD4sp T cells that removes the inconsistencies and safety issues caused by using xenogenic materials. iPSCs can serve as an endless supply of CD4sp T cells and Tregs with and without a CAR for therapeutic applications and our study supports further development of this process for clinical use in oncology and autoimmune diseases.

## Limitations of the study

Further characterization of the iPSC-Tregs and iPSC-CD4sp T cells is still required, including evaluation ability to expand to clinically relevant numbers and *in vivo* functions. Our initial transcriptomics data shows that the iPSC-CD4sp T cells generated have an immature and innate-like phenotype. Further optimization of the iPSC-CD4sp differentiation can be achieved using a model-guided approach such as design of experiment (DOE) to optimize culture conditions and a more in-depth multi-omics analysis to identify genes that can be modified through genome engineering to obtain a more mature CD4 T cell phenotype. While we have clearly shown that the iPSC-Tregs are functional *in vitro*, it is essential to test their suppressive capacities and phenotypic stability *in vivo* in a GvHD mouse model.

## Methods

### Generation of human induced pluripotent stem cell lines

Previously cryopreserved CD4 T cells were used for reprogramming to pluripotency. CD4 T cells were isolated from mobilized leukapheresis material from a healthy male donor (AllCells, LLC) with informed written consent. The raw material was washed several times to remove platelets, labeled with CD4 microbeads (Miltenyi Biotec) and then isolated using the Miltenyi CliniMACS system. Isolated CD4 T cells were analyzed for purity prior to cryopreservation. To prepare for reprogramming, the CD4 T cells were thawed and cultured overnight in RPMI + 10% human AB serum (huABS, Valley Biomedical) and activated 1:1 with CD3/CD28 Dynabeads in 100 U/mL of IL-2 (Thermofisher). The next day, the CD4 T cells were debeaded and sorted for CD4^+^CD25^+^ cells and recovered overnight in RPMI + 10% huABS with 100 U/mL of IL-2. One day post-sort, the sorted CD4 T cells were reprogrammed with Sendai virus carrying the four transcription factors OCT, SOX2, KLF4, and C-MYC as previously reported using Cytotune iPS-2.0 Sendai Reprogramming Kit (Thermofisher; Fusaki Proc Japan Academy 2008). After transduction, cells were recovered in RPMI + 10% huABS for 24 hours and transferred to ReproTeSR media (StemCell Technologies) on Matrigel-coated plates (Corning). Cells were fed daily or every day until the appearance of colonies by day 14. Colonies were examined for appropriate morphology then manually selected for further expansion and transition to mTeSR1 media (StemCell Technologies)

### Characterization of human induced pluripotent cells lines

Immunocytochemical analysis for pluripotency of iPSC lines was performed by washing cells in 1X PBS (Thermofisher), fixed in 4% paraformaldehyde for 20 minutes at 4°C, and blocked in 10% normal goat or rabbit serum with 0.5% Triton X-100 (Sigma) in PBS. Immunocytochemical analysis was performed using primary antibodies and corresponding AlexaFluor-conjugated secondary antibodies (Thermofisher). All antibodies were diluted and used according to manufacturer’s instructions. Cell nuclei were counterstained with DAPI in some experiments^72^. Images were collected using an EVOS Cell Imaging System (Thermofisher). PluriTest analysis was performed by ThermoFisher Scientific. Karyotype analyses were performed by Cell Line Genetics.

### iPSC culture and clonal expansion

iPSCs were maintained on Matrigel (Corning) in mTeSR1 media according to manufacturer’s instructions (StemCell Technologies). For passaging, iPSC colonies were detached briefly with either Accutase and manually scraped or with Gentle Cell Dissociation Reagent. Cells were routinely passaged at a 1:8-1:10 split ratio in the presence of Y-27632 (ROCK inhibitor, Tocris) for the first day after passaging and removed for subsequent media changes.

For clonal expansion of transfected iPSCs, a method of low-density passaging was used. After electroporation, iPSCs were passaged at a low seeding density (1e3, 2.5e3, 5e3 cells per well of a 6-well plate) such that a single iPSC would give rise to a single colony in a Matrigel-coated 6-well plate. iPSC colonies were then manually picked and expanded individually. Tightly packed colonies with clearly defined borders were then collected for genomic DNA isolation and subsequent screening for targeted integration.

### Differentiation of HSPCs from iPSCs

iPSCs were differentiated as previously described with modifications^24, 25^. Embryoid bodies were generated from iPSCs in APEL2 media (StemCell Technologies) and cultured in non-tissue culture or ultra-low attachment 96-well round bottom plates in the presence of 10 ng/mL BMP4, 10 ng/mL VEGF, 50 ng/mL SCF, 10 ng/mL bFGF, and 10 μM Y-27632 on day 0 (all cytokines from Thermofisher). EBs were then cultured in APEL2 media in the presence of 20 ng/mL VEGF, 10 ng/mL bFGF, 100 ng/mL SCF, 20 ng/mL FLT-3L, 20 ng/mL TPO and 40 ng/mL IL-3 from day 1 to day 6 with a media refresh every 2 days (all cytokines from Thermofisher). On day 8, EBs were switched to complete StemPro-34 media (Thermofisher) and supplemented with 100 ng/mL SCF, 20 ng/mL FLT3-L, 20 ng/mL TPO, and 40 ng/mL IL-3 in order to expand developing HSPCs. Media was refreshed every 2 days until day 14. The EBs were then collected and passed through a 70 μM cell strainer. The resulting cell suspension containing the HSPCs was then used for downstream differentiation.

### Differentiation of T cell progenitors from iPSC-HSPCs

StemDiff T cell kit (StemCell Technologies) was used to differentiate HSPCs to lymphoid progenitor cells and to DP T cells. Manufacturer’s instructions were followed unless otherwise indicated. HSPCs were seeded at 50,000 cells/well of a 6-well plate coated with lymphocyte differentiation coating material (LDCM) and cultured in StemSpan Lymphoid Progenitor Expansion Media to differentiate the HSPCs to CD5+CD7+ lymphoid progenitor cells. After 14 days of culture (differentiation day 28), resulting iPSC-lymphoid progenitor cells were harvested and seeded at 0.5e6/well of a 6-well plate with a fresh coating of LDCM. The cells were cultured in StemSpan T Cell Progenitor Maturation Media (maturation media) for differentiation to CD4+CD8+ DP T cells. After 14 days of culture (differentiation day 42), the DP T cells were harvested for further differentiation.

### Generation of CD4-single positive T cells and Tregs from iPSC-DP T cells

Induction of DP T cells to CD4sp T cells was performed on LDCM in maturation media. On differentiation day 42, a cell stimulation cocktail composed of phorbol 12-myristate 13-acetate and ionomycin was added to the cultures at three different concentrations from the 500X stock concentration: low (0.01 μM PMA, 0.16 μM Ionomycin), medium (0.02 PMA μM , 0.335 μM Ionomycin), and high (0.0405 μM PMA, 0.67 μM Ionomycin). A full media change was performed 24 hours later by harvesting the entire cell suspension, replating into fresh maturation media and seeding at a 1:1 ratio onto new LDCM coated 6-well plates. The cells were then fed every 3-4 days for an additional week until day 50. On day 50, the cells were then treated with 20 uL/mL of Immunocult Human Treg Differentiation Supplement (StemCell Technologies) containing recombinant human TGF-B1 and all-trans retinoic acid, 12.5 uL/mL of Immunocult Human CD3/CD28/CD2 T cell activator (StemCell Technologies) and 10 ng/mL of CTS IL-2 (Thermofisher). Media in the wells for conversion to Tregs was composed of a 50% mixture of complete CTS OpTmizer and maturation media. A half media change was performed on day 54 with a 50% mixture of CTS OpTmizer and maturation media supplemented with 10 ng/mL of IL-2.

### Flow cytometry, sorting and antibodies

Flow cytometry was performed in FACS buffer composed of PBS, 0.5% BSA, and 1 mM EDTA. For extracellular staining, antibodies were added to cells according to manufacturer’s instructions, incubated at 4°C for 20 minutes, and washed before analysis. For intracellular staining, cells were first stained with extracellular markers and fixed and permeabilized using the eBioscience FOXP3/Transcription Factor Staining Buffer set (Thermofisher). Antibodies were then added according to manufacturer’s instructions. Attune NxT flow cytometer (Thermofisher) and Sony Sorter SH800 was used for flow cytometric analysis and sorting, respectively. Antibodies (clones in parentheses) used for surface and intracellular staining of the following molecules were obtained from Biolegend: CD45-FITC (Clone HI30, #304054), FOXP3-AF647 (Clone 206D, #320114), ThPOK-PE (Clone 11H11A14, #656407), TCRab-PECy (Clone IP26, #306720), Helios-PE (Clone 22F6, Cat# 137216), CD127-BV605 (Clone A019D5, #351334), CD127-AF700 (Clone A019D5, #351344), CD28-FITC (Clone S20013F, #377612), CD62L-PECy7 (Clone MEL-14, #104418), and PD-1-APC (Clone 12.2H7, #329908). Antibodies against CD34-APC (Clone 581, #555824), CD25-PECy7 (Clone 2A3, #335789), CD45RO-BV421 (Clone UCHL1, #562641) and CD3-BV510 (Clone UCHT1, #563109) were obtained from BD Biosciences. Antibodies against CD4-e450 (Clone OKT-4, #48-0047-42), CD4-PE (Clone OKT-4, #12-0048-42), CD8-FITC (Clone SK1, #11-0087-42), and CD8-APC (Clone SK1, #11-0087-42) were obtained from Thermofisher. CD69-APC (Clone REA824, #130-113-219), CD5-PE (Clone UCHT1, #130-119-855) and CD7-APC (Clone CD7-687, #130-123-265) antibodies were obtained from Miltenyi. An antibody to CD25-PE (Clone 2A3, #60153PE) was obtained from StemCell Technologies. Antibodies to SOX2 (#MAB2018), OCT4 (#AF1759), SSEA-4 (#MAB1435) were obtained from R&D Systems. TRA-1–81 (#MAB8495) was obtained from Santa Cruz Biotechnology. The following fluorophore-conjugated secondary antibodies and DAPI were obtained from Thermofisher: donkey-anti-mouse secondary antibody Alexa-Fluor 594 (#A21203) and donkey-anti-goat secondary antibody Alexa-Fluor 488 (#A11055).

### scRNA-sequencing RNA extraction, library generation, and sequencing analysis

RNA from iPSC-Tregs, iPSC-CD4sp, primary Tregs, induced Tregs and primary Tconv was extracted using an RNA extraction kit (Qiagen) following differentiation (iPSC-Tregs and induced Tregs) or isolation and culture (primary Tregs and Tconv). Primary cells were isolated from leukapheresis. Leukapheresis were obtained from healthy volunteers who provided written informed consent.

Single cell RNA-sequencing libraries were sequenced on a NextSeq 500. Samples had an average depth of 18636 reads per cell. Read quality control, UMI counting, barcode counting, and alignment to the GRCh38 GENCODE v32/Ensembl 98 reference genome from 10x Genomics were performed using the “cellranger 6.0.2” pipeline. All downstream analysis were performed using the Seurat packages in R for single-cell RNAseq analysis^73^.

All samples were filtered on the following measures: number of unique genes detected, total number of molecules detected, and level of expression of mitochondrial reads before integrating the datasets. We removed cells in which greater than 20% mitochondrial reads, less than 200 feature counts, or greater than 6500 feature counts were detected.

The data was then normalized using the Seurat normalization function which divides the feature counts by total counts for each cell, then multiplies the resulting fraction by a scale factor of 1.00e+4, and, lastly, log-normalizes this value.

Dimensionality reduction was performed using Seurat packages and utilizing principal components 1 through 50 in the reduction. Clusters were determined at a resolution of 0.3. The identity of each cluster was validated by evaluation of gene expression within each cluster of cell-type specific markers for the cell-types of interest.

The Seurat object was exported and converted to an h5ad file to use CellTypist’s automated cell type annotation. CellTypist’s Immune_All_High, Immune_All_Low, and Pan_Fetal_Human models were used for the automated annotation of cell types^74^. The mode “prob match” was used for the Pan_Fetal_Human model to assign multiple cell types per sample.

### Assessment of iPSC-CD4sp T cell and iPSC-Treg Function

For assessment of Tconv or iPSC-CD4sp T cell phenotype and activation, cells were sorted and cultured in X-VIVO15 (Lonza) + 10% huABS (Valley Biomedical). Cells were either left unactivated or activated with Immunocult Human CD3/CD28/CD2 T cell activator per manufacturer’s instructions (StemCell Technologies) for 24 or 48hrs. Cells were then stained with indicated antibodies and run on flow cytometry as previously described. For assessment of cytokine secretion, unactivated or activated iPSC-CD4sp T cells were seeded at 5e4 cells/well in a round-bottom 96 well plate in 200 uL of media. The supernatant was collected 24 hours later.

For assessment of iPSC-Treg function, cells were sorted on day 57 and cultured in complete OpTmizer media supplemented with 1000U IL-2/mL (Thermofisher) for three days prior to functional analysis. iPSC-Tregs were then activated with CD3/CD28 Dynabeads (Thermofisher) at a ratio of 1:1 in media composed of X-VIVO15 (Lonza) + 10% huABS (Valley Biomedical) + Penicillin / Streptomycin solution (pen/strep). For analysis of cytokine production, 5e4 activated and unactivated cells were seeded in a round-bottom 96-well plate in 200 uL of media. One day post-activation, supernatant was collected and analyzed for cytokines using human cytokine assay X (Meso Scale Discovery). For assessment of Tconv suppression, activated and unactivated iPSC-Tregs were co-cultured with activated (2:1 beads:cells) Tconv labeled with Cell Trace Violet (Thermofisher) in X-VIVO15 media supplemented with pen/strep. iPSC-Tregs were initially plated at 5e4/well in a 96-well round-bottom plate and serially diluted by a factor of 2 down to 781.25 cells/well. 2.5e4 activated, CTV-labeled Tconv were then added to each serially diluted well of iPSC-Tregs. Baseline Tconv proliferation was determined by culturing 2.5e4 Tconv in the absence of iPSC-Tregs. Three days after co-culture, cells were collected, labeled for viability and assessed for Tconv proliferation via Cell Trace Violet dye by flow cytometric analysis. For assessment of iPSC-Treg cytolytic activity, activated and unactivated iPSC-Tregs were cocultured with activated Tconv. 5e4 iPSC-Tregs were seeded with 5e4 primary Tconv and cultured for 3 days in X-VIVO15 media supplemented with pen/strep. The supernatant was then collected and analyzed using the CyQuant LDH Cytotoxicity Kit (Thermofisher) following the manufacturer’s instructions. Samples were read on a SpectraMax microplate reader (Molecular Devices). Primary Treg, Tconv and CD8 controls were cultured for 7-9 days prior to activation and assessment of function.

### FOXP3 TSDR demethylation assay

The Treg-specific demethylated region (TSDR) hypomethylation status of FOXP3 gene was measured using a post-PCR High Resolution Melt (HRM) analysis as previously described^6^. Briefly, genomic DNA was extracted from cell pellets containing at least 1e6 cells/pellet using the DNeasy Blood and tissue kit (Qiagen), followed by bisulfite conversion using the Epitech Fast Bisulfite Conversion Kit (Qiagen). The FOXP3 TSDR regions were then amplified from the converted sample gDNAs by qPCR using specific primers (forward: TTGGGTTAAGTTTGTTGTAGGATAG, reverse: ATCTAAACCCTATTATCACAACCCC, Sigma Aldrich, USA) and Precision Melt Supermix (Biorad). The melting curves of amplified product DNA fragments were quantified against the methylation standard curve generated using commercial bisulfite converted human genomic DNAs (Qiagen) and expressed as a mean methylation percentage.

### Cytokine MSD methods

The concentrations of cytokines were measured with MSD U-Plex kits (#K15067L-1, MSD, USA) following the manufacturer’s protocol. Briefly, biotinylated capture antibodies were incubated with specific linkers and then the mixture of linker coupled antibodies was added to MSD U-Plex plates. After 1 hour incubation and wash step, standard curves and samples were added to the plates. The plates were washed again after the sample incubation. A mixture of SULFO-Tag conjugated detection antibodies was added to the wells and incubated. After the final wash step, MSD GOLD Read Buffer B was added to the plate followed by reading the electrochemiluminescence signals in a MESO SECTOR S 600 instrument.

### ZFN and donor DNA constructs

The zinc finger nuclease (ZFN) pair targets the *TRAC* locus, encoding for the alpha chain of the TCR, at exon 2. The ZFP backbone and FokI variants were generated by reassembling zinc finger arrays from modules as previously described ^75^ and cloning the resulting zinc finger array into a vector containing the desired FokI variant. All constructs were verified by Sanger sequencing with two sequencing primers to yield overlapping sequence reads of the entire ZFN coding sequence^76^. mRNA was transcribed from a PCR template using the 5×m Message mMachine T7 ULTRA kit (Invitrogen) according to the manufacturer’s instructions as previously reported^76^.

We designed and cloned the donor DNA sequence into a customized AAV vector containing AAV inverted terminal repeats (ITRs). An AAV vector was initially selected to use AAV as DNA delivery system but we ultimately optimized delivery using plasmid DNA. The pAAV-partial-TRAC-HLA-A2 CAR contains 110bp of genomic partial TRAC sequence (amplified by PCR) flanking the TRAC homologous donor templates in frame with the second exon of TRAC, followed by a self-cleaving T2A peptide, the HLA-A2 CAR and a bGHpA sequence.

### Transfection of TRAC-ZFN mRNA and donor DNA

All transfections were performed on the Maxcyte ATx (model GT) electroporation system. Approximately 3e6 iPSCs were used for electroporation in an OC-100 processing assembly (Maxcyte). iPSCs were treated with Accutase until cells were fully detached (∼3min). After collection, cells were pelleted and washed once with Maxcyte electroporation buffer. The iPSCs were then resuspended in Maxcyte electroporation buffer at a concentration of 30e6 cells/mL and mixed with ZFN mRNA (10 ng/μl) and 8 ug of donor DNA. The cell and nucleic acid mixture was dispensed into the OC-100 assembly and immediately placed into the Maxcyte instrument using Optimization program 9. iPSCs were then recovered for 20 minutes at 37°C in the assembly and then transferred to Matrigel-coated plates in mTeSR1 media. Y-27632 was added to the media for 2-3 days post-transduction until confluent and then passaged for clonal expansion.

### Analysis of targeted integration in iPSCs

iPSCs were assessed quantitatively for on-target insertions/deletions and targeted integration through NGS sequencing on the Illumina platform. Genomic DNA from clonally expanded iPSCs was first amplified, followed by nested PCR amplification to prepare ∼200 bp amplicons for paired-end deep sequencing on the Illumina MiSeq sequencer.

### iPSC-Treg Cryopreservation

Post-sort CAR iPSC-Tregs were recovered in complete OpTmizer supplemented with 1000U IL-2/mL at a concentration of 1e6/mL. 24 hours after recovery, CAR iPS-Tregs were harvested, counted and cryopreserved in CryoStor CS10 (Biolife Solutions) at a concentration of 4-5e6/mL. Cryopreserved cells were placed in an alcohol-free freezing container (CoolCell, Corning) and placed at -80C overnight before transferring to LN2 for long-term storage.

## Data Availability

Data will be made available upon reasonable request. Sequencing data to prove targeted insertion and scRNAseq data will be deposited on relevant NCBI archives before publication and access codes will be shared. Due to the proprietary nature of the donor plasmid sequence we have restricted data access to the full-length sequencing data of the insert for the edited iPSC cell lines as described in the manuscript.

## Material Availability

For this research we have derived unique iPSC lines (edited and non-edited) from human T cells and developed proprietary Zinc finger nucleases (ZFN). We are restricting access to these lines and ZFN sequences and mRNA to protect Sangamo Therapeutics’ proprietary work. Any other requested materials will be reviewed on a case-by-case base and be subject to a Material Transfer Agreement.

## Ethics Statement

The leukapheresis collection at AllCells, LLC used for the derivation of the iPSC cell line was approved by an independent review board (Alpha IRB) under the study title “Non-Mobilized Mononuclear Cell Apheresis Collection from Healthy Donors for the Research Market” (Protocol number: 7000-SOP-045). Informed consent was obtained from the cell donor. Specially trained nurses at LeukoLab, a division of AllCells, collected the leukopack from the donor. The donor was compensated for their time and effort related to their participation in the study as approved by the IRB.

## Acknowledgements

We thank Prof. Maria Themeli for critical review of the manuscript and for giving critical advice for our research work. We also thank Prof. Megan Levings for critical advice on the transcriptomics analysis; Dr. Andreas Reik, Dr. Lei Zhang, Dr. George Kwong and Dr. Steven Cincotta for technical assistance; Dr. Julie Gertner-Dardenne, Dr. Maurus de la Rosa, Dr. David Fenard, Dr. Tobias Abel and Dr. Virginie Fabre for scientific discussions; Cayla McEwen and Jean Alladin for administrative support.

## Patents

Part of this research work is described in the following patents:

US Patent number 11512287: “Targeted disruption of T cell and/or HLA receptors”

Application PCT/US2021/057152 (Filed Oct 28, 2021): “Generation of CD4+ Effector and Regulatory T cells from Human Pluripotent Stem Cells”

Application PCT/US2020/059730 (Filed Sep 11, 2020): “Generation of Engineered Regulatory T cells”

## Author contributions

H.F., M.M., L.N., G.L., S.G., A.C. and S.Z. conception and design; H.F., M.M., J.J., L.N., K.K., S.S., J.H. and A.V. collection and/or assembly of data; H.F., M.M., J.J., L.N., K.K., G.L., S.G., S.S., J.H. and A.V. data analysis; H.F., M.M., L.N., S.S., A.C., J.F. and S.Z. data interpretation; J.F. advice on experimental design; A.C. and S.Z. study supervision; H.F. and S.Z. manuscript writing.

