## Supplemental Figures File for "A Serum- and Feeder-Free System to Generate CD4 and Regulatory T Cells from Human iPSCs"

**SUPPLEMENTARY FIGURES**

**SUPPLEMENTARY FIGURE 1. Characterization of iPSCs**

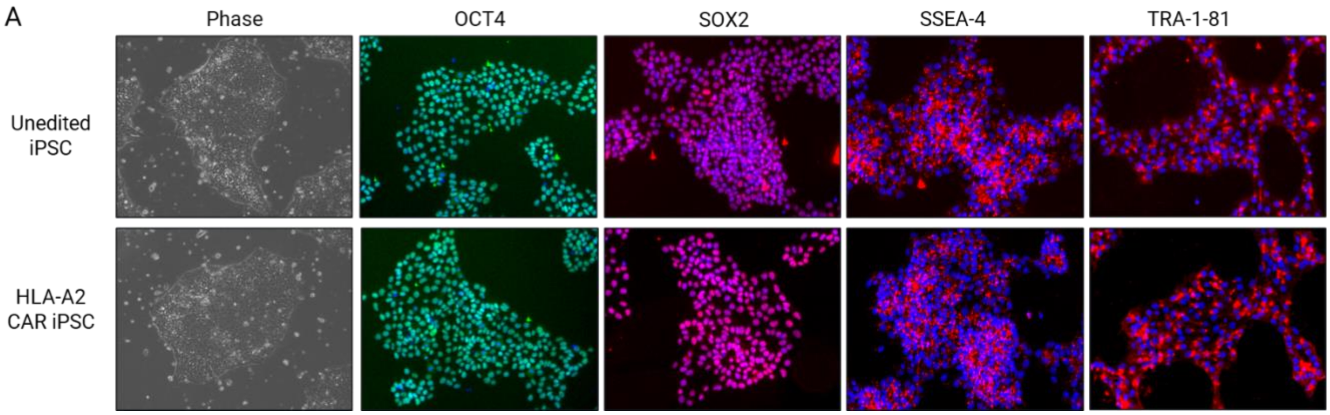


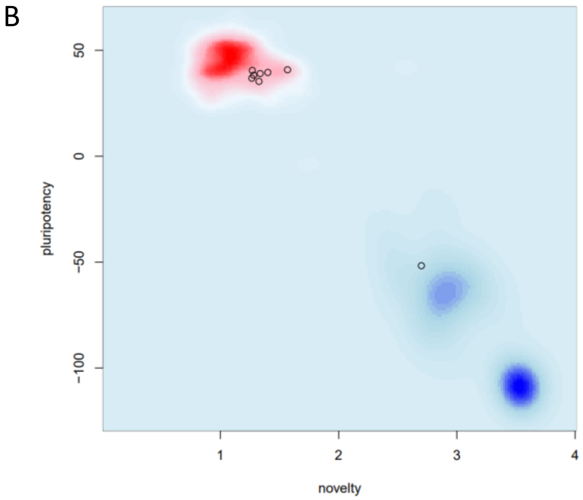


| Sample | Sample ID | Result | PluriCor | NovelCor |
|---|---|---|---|---|
| PT-5040 | iPSC unedited line 1 | Pass | 38.23591 | 1.282135 |
| PT-5041 | iPSC unedited line 2 | Pass | 36.83992 | 1.265971 |
| PT-5042 | iPSC unedited line 3 | Pass | 39.53625 | 1.401372 |
| PT-5043 | iPSC HLA-A2 CAR line 1 | Pass | 40.79364 | 1.567719 |
| PT-5044 | iPSC HLA-A2 CAR line 2 | Pass | 35.33185 | 1.326469 |
| PT-5045 | iPSC HLA-A2 CAR line 3 | Pass | 39.02393 | 1.336279 |
| PT-5055 | iPSC Control | Pass | 40.50378 | 1.270098 |
| PT-5056 | Non iPSC Control | Fail | -5168477 | 2.700077 |

C

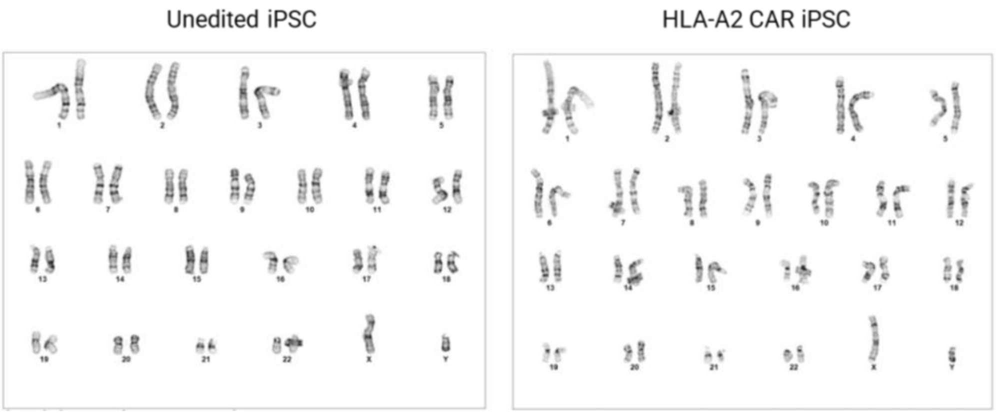


**Supp. Figure 1. Characterization of iPSCs.**

A. Representative images of T cell-derived iPSCs and HLA-A2 CAR edited iPSCs. iPSCs were assessed for morphology by phase contrast microscopy and expression of OCT4, SOX2, SSEA-4, and TRA-1-81 by immunocytochemistry.
B. PluriTest results of T cell-derived iPSCs and HLA-A2-targeted CAR edited iPSCs confirm pluripotency. Samples were compared to positive and negative control cell lines.
C. Representative G-band karyotyping analyses confirming normal 46 XY karyotype in T cell-derived iPSCs (3 clones) and HLA-A2-targeted CAR-edited iPSCs (3 clones).

**SUPPLEMENTARY FIGURE 2. Double positive (DP) T Cell Yield from Unedited and HLA-A2 CAR iPSCs with CXCL12 and p38i Treatment.**

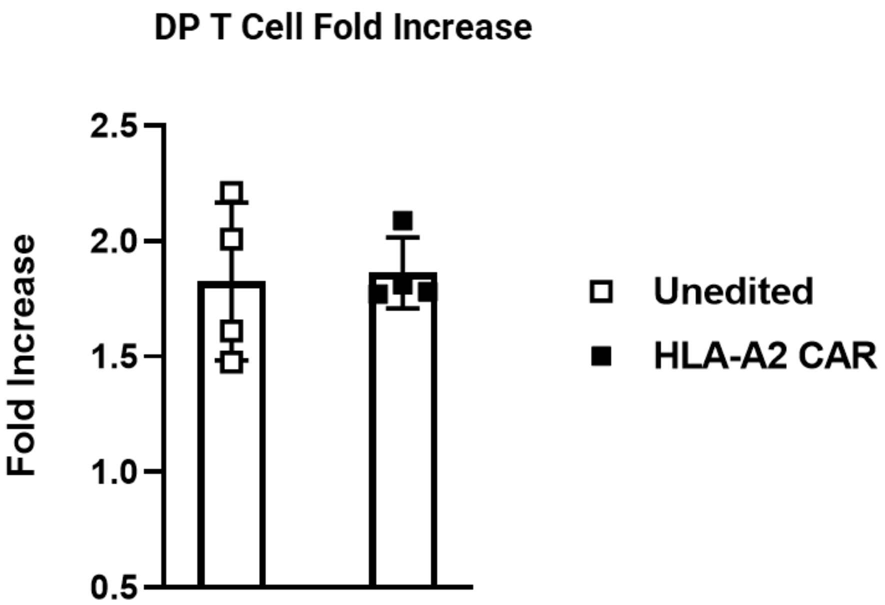


**Supp. Figure 2. Double positive (DP) T cell yield from Unedited and HLA-A2 CAR iPSCs with CXCL12 and p38i treatment.**
Yield of DP T cells from unedited and CAR edited iPSCs after 42 days of differentiation. CXCL12 and p38i were added at day 28 of differentiation (n=4 from n=1 clone, unedited; n=4 from n=1 clone, HLA-A2 CAR). Data represent mean ± SD of n independent experiments.

**SUPPLEMENTARY FIGURE 3. Induction of iPSC-CD4sp T cells by PMA+I**

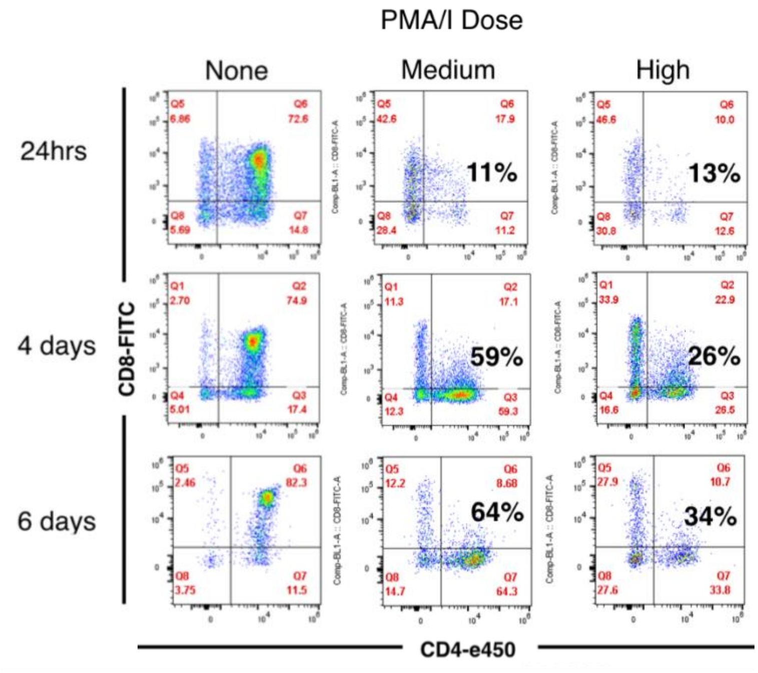


**Supp. Figure 3. Induction of iPSC-CD4sp T cells by PMA+I.**

Representative kinetic analysis of developing CD4sp T cells after Phorbol 12-myristate 13-acetate and Ionomycin (PMA/I) addition (medium and high dose). Phenotype was assessed by the expression levels of CD4 and CD8 by flow cytometry (n=1).

**SUPPLEMENTARY FIGURE 4. iPSC-CD4sp phenotype**

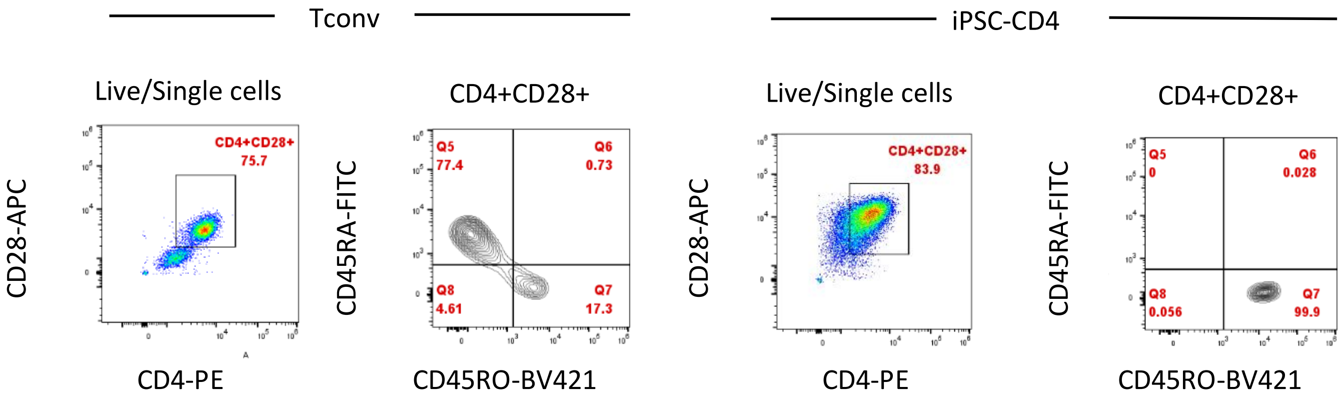


**Supp. Figure 4. iPSC-CD4sp T phenotype.**

Representative flow plots and expression of CD4 markers in unactivated T conventional (Tconv) and sorted unactivated iPSC-CD4sp T cells. Phenotype was assessed by flow cytometric analysis in Tconv (n=3) and iPSC-CD4sp (n=3 from n=1 clone) at day 50 of differentiation for the expression of CD4, CD62L, CD45RO and CD28.

**SUPPLEMENTARY FIGURE 5. Sorting Strategy for iPSC-CD4sp and for iPSC-Tregs**

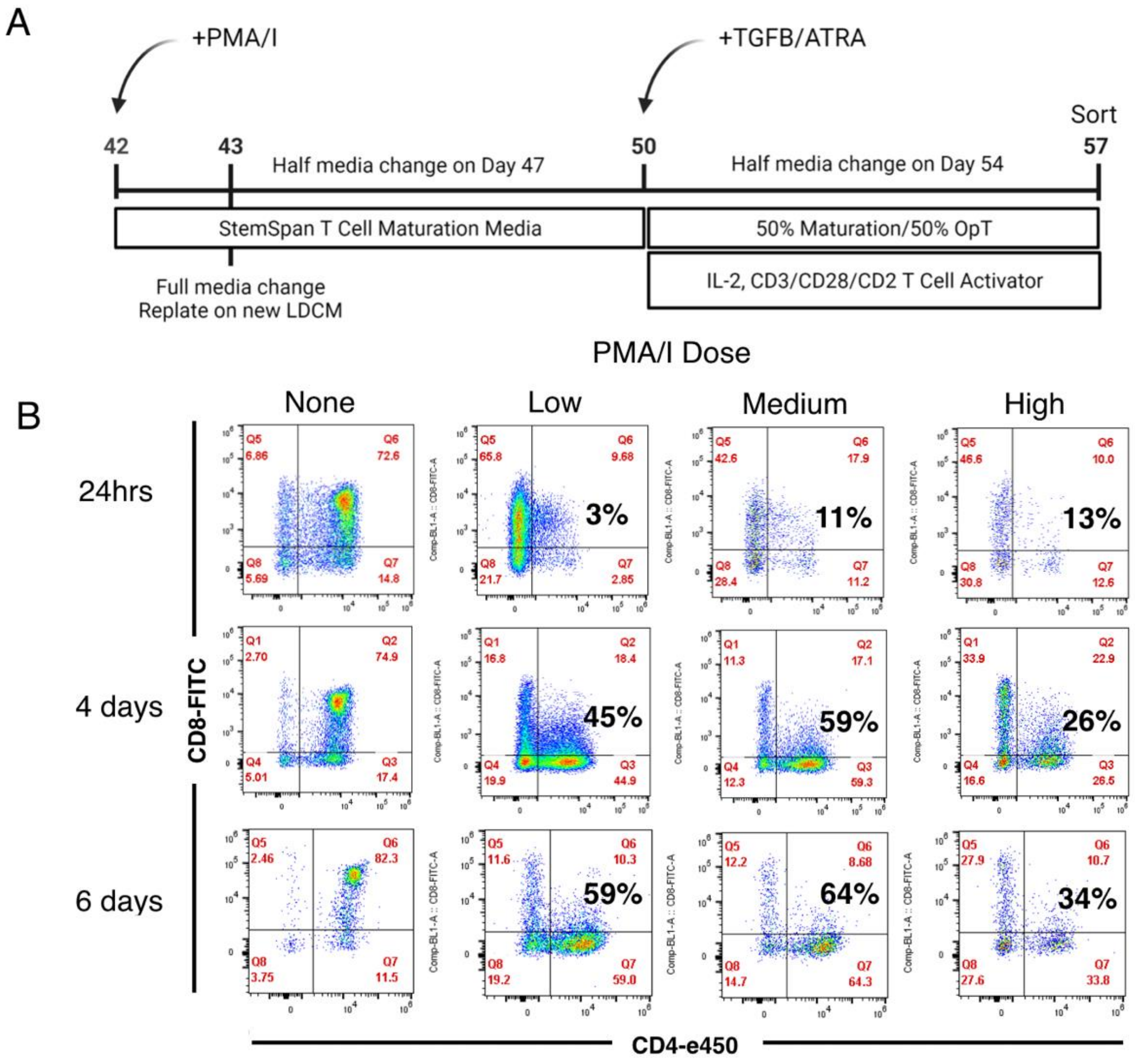


**Supp. Figure 5. Sorting strategy for iPSC-CD4sp T cells and iPSC-Tregs.**

Representative flow plots of iPSC-Treg T cell sorting at day 57. The same CD4 gate was used to sort iPSC-CD4sp T cells at day 50.

**SUPPLEMENTARY FIGURE 6. Automated cell type annotation of scRNA-seq data.**

A

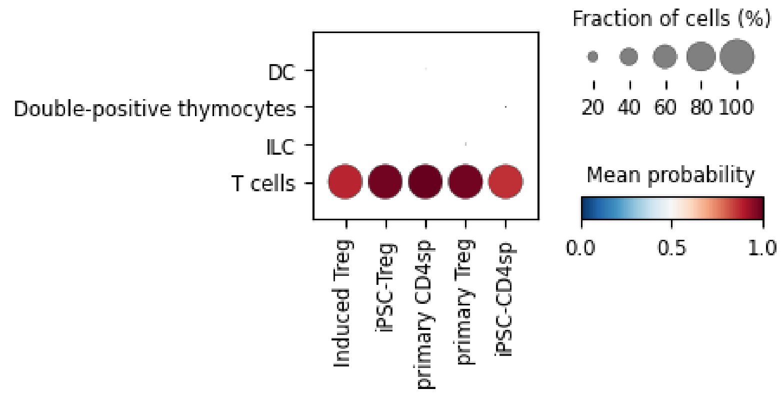


B

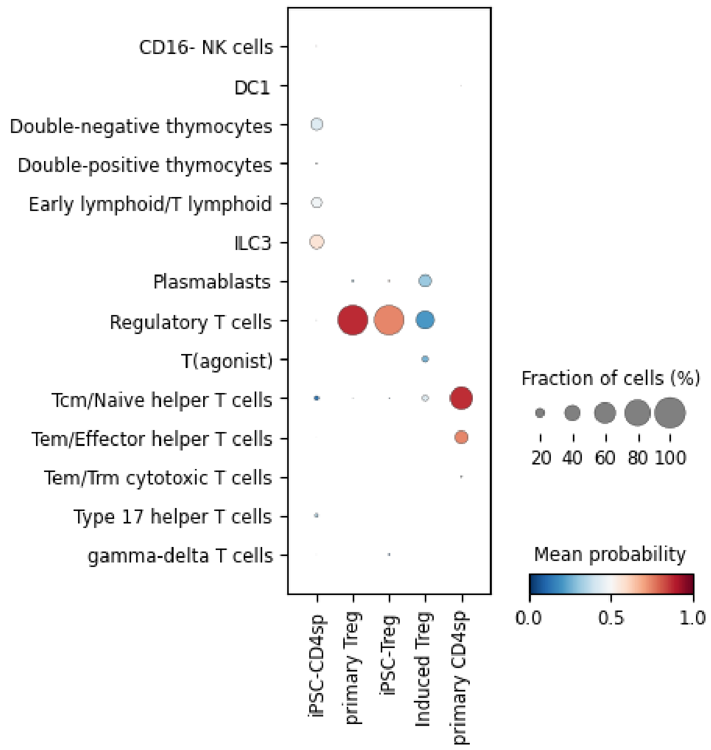

C

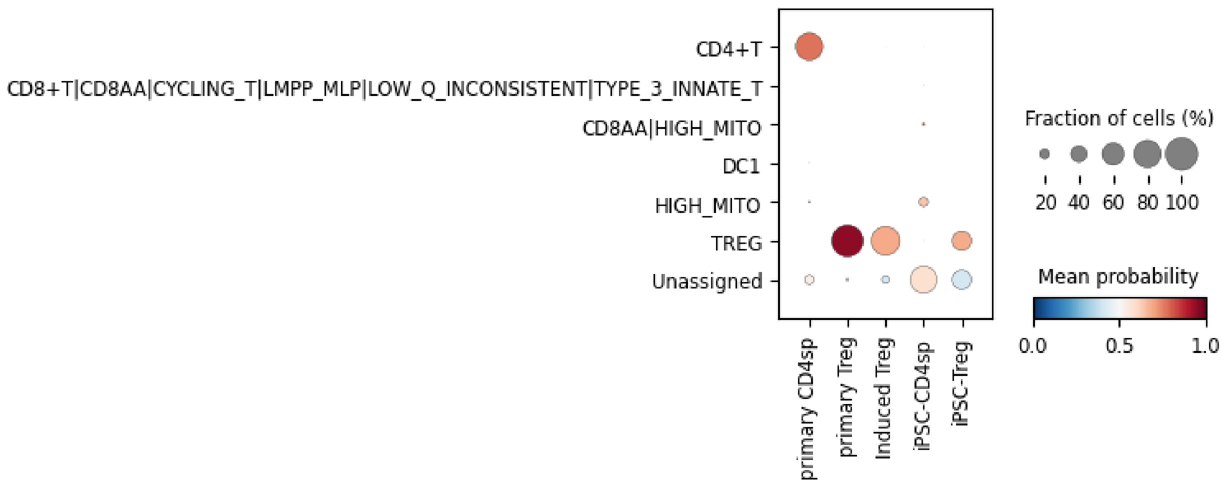
CD4+T
CD8+T|CD8AA|CYCLING_T|LMPP_MLP|LOW_Q_INCONSISTENT|TYPE_3_INNATE_T
CD8AA|HIGH_MITO
DC1
HIGH_MITO
TREG
Unassigned
primary CD4sp
primary Treg
Induced Treg
iPSC-CD4sp
iPSC-Treg
Fraction of cells (%)
20 40 60 80 100
Mean probability
0.0 0.5 1.0

D

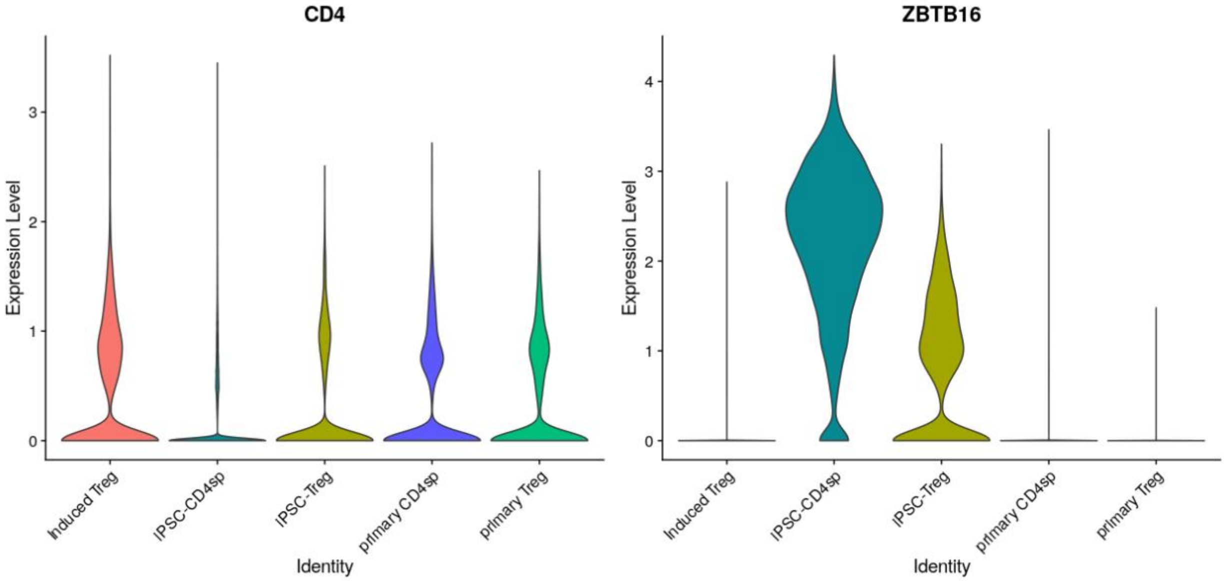
CD4
ZBTB16
Expression Level
Identity
Induced Treg
iPSC-CD4sp
iPSC-Treg
primary CD4sp
primary Treg

**Supp. Figure 6. Automated cell type annotation of scRNA-seq data.**

A. Dot plot comparing the prediction result of CellTypist against the known cell types of induced Treg, iPSC-Treg, primary CD4sp, primary Treg and primary CD4sp using the Immune_All_High model (adult immune cells).
B. Dot plot comparing the prediction result of CellTypist against the known cell types of induced Treg, iPSC-Treg, primary CD4sp, primary Treg and primary CD4sp using the Immune_All_Low model (adult immune cell types).
C. Dot plot comparing the prediction results of CellTypist against the know cell types of induced Treg, iPSC-Treg, primary CD4sp, primary Treg and primary CD4sp using the Pan_Fetal_Immune model (fetal immune cell types) allowing for multiple cell type classifications.
D. Violin plot for *CD4* and *ZBTB16* expression across cell types.

**SUPPLEMENTARY FIGURE 7. Characterization of sorted and activated iPSC-CD4sp.**

A

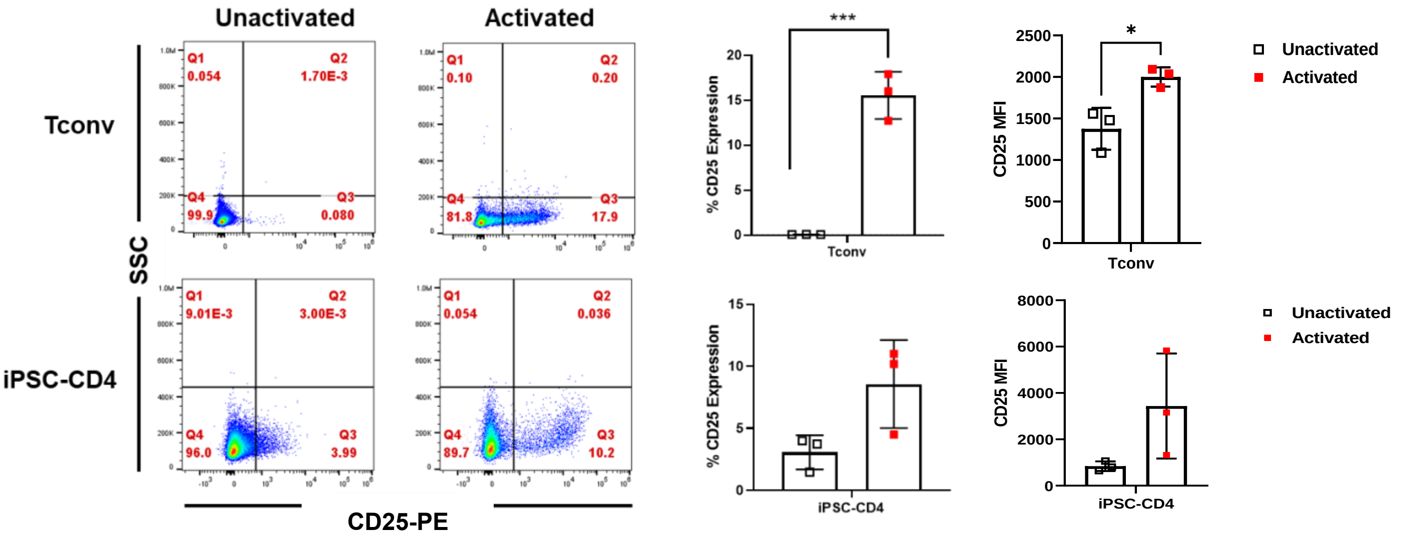


B

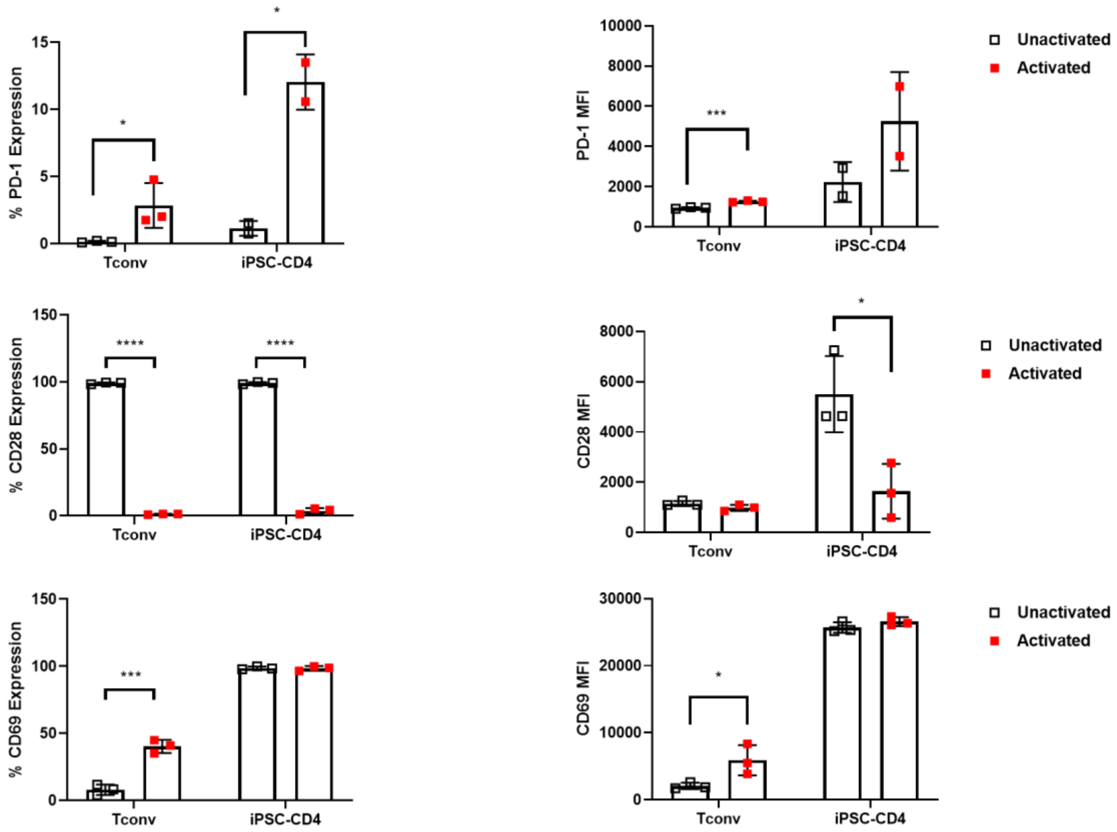

**Supp. Figure 7. iPSC-CD4sp T cell characterization activation markers at 24 hours post-stimulation**

A. Representative flow plots and expression of CD25 in sorted unactivated and activated iPSC-CD4sp T cells or T conventional (Tconv) cells (n=3 from n=1 clone) by flow cytometry. CD25 Median fluorescence intensity (MFI) was calculated (right).
B. Expression of activation markers (CD69, PD-1 and CD28) was assessed by flow cytometry after 24 hours of activation. MFI was calculated (right).
   Data show either a representative flow plot set of n independent experiments or represent mean ± SD of n independent experiments. P*<0.05, P***<0.001, P****<0.0001 by unpaired two-tailed t-test.

**SUPPLEMENTARY FIGURE 8. Characterization of HLA-A2 CAR+ iPSC-Tregs pre-sort**

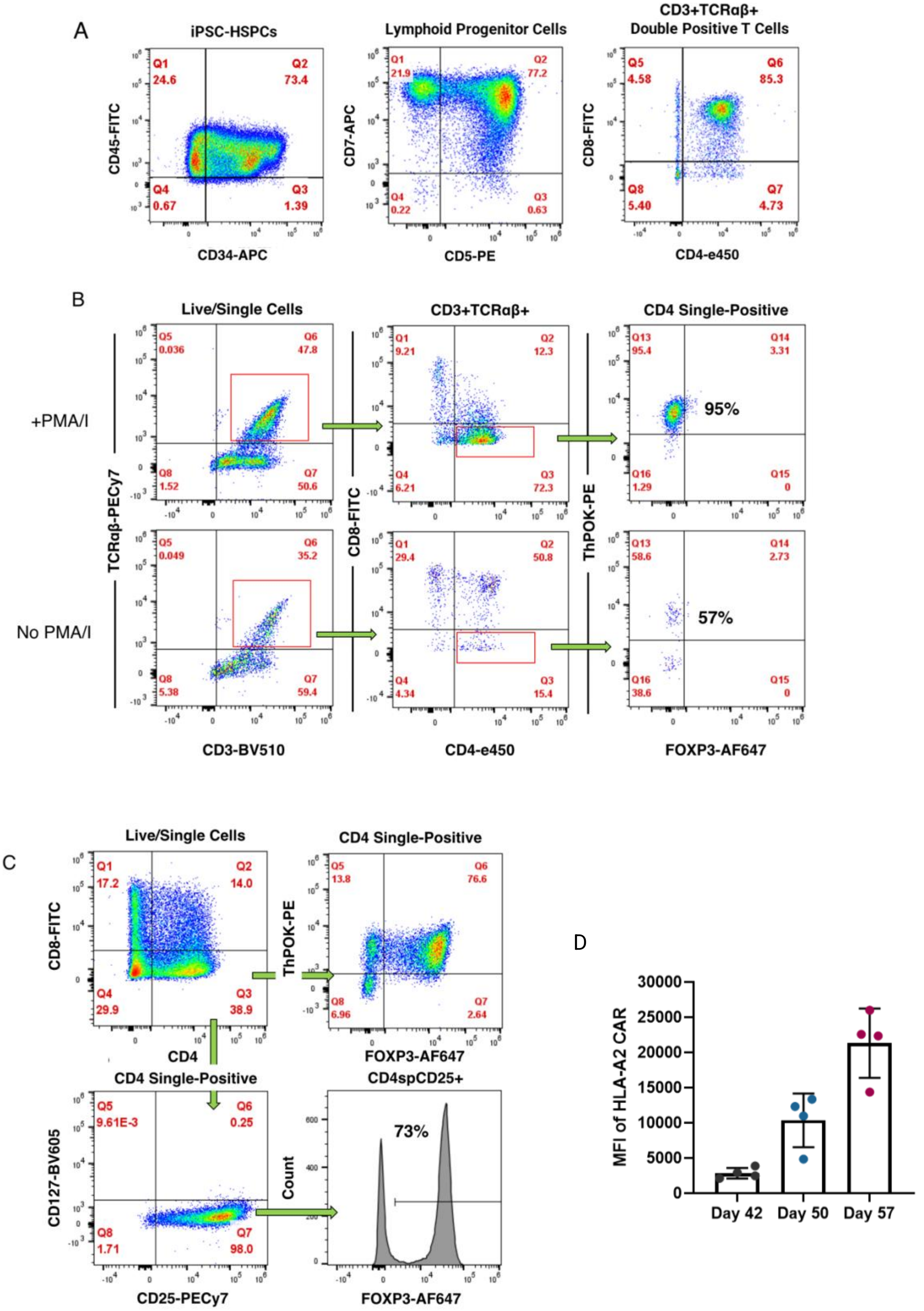

**Supp. Figure 8. Characterization of HLA-A2 CAR+ iPSC-Tregs**

A. Representative flow cytometric plots of hematopoietic stem and progenitor cells (HSPCs), lymphoid progenitor cells, and double positive (DP) T cells from HLA-A2 CAR edited iPSCs (n=3 from n=3 clones).
B. Representative flow plots of iPSC-CD4sp T cells 8 days after PMA/I addition (day 50 of differentiation). Phenotype was assessed by the expression levels of CD3, TCRαβ, CD4, CD8, FOXP3 and ThPOK in cells with and without PMA/I treatment (n=3 from n=3 clones).
C. Representative flow plots of iPSC-Tregs 7 days after TGFβ+ATRA addition (differentiation day 50). CD4sp phenotype was assessed by the expression levels of CD3, TCRαβ, CD4, CD8, FOXP3 and ThPOK (n=3 from n=3 clones). Treg phenotype was assessed by the expression levels of CD4, CD8, CD25, CD127, and FOXP3 (n=3 from n=3 clones).
D. MFI of CAR expression as measured at differentiation days 42, 50, and 57.
Data represent mean ± SD of n independent experiments.

**SUPPLEMENTARY FIGURE 9. Characterization of sorted HLA-A2 CAR+ iPSC-Tregs**

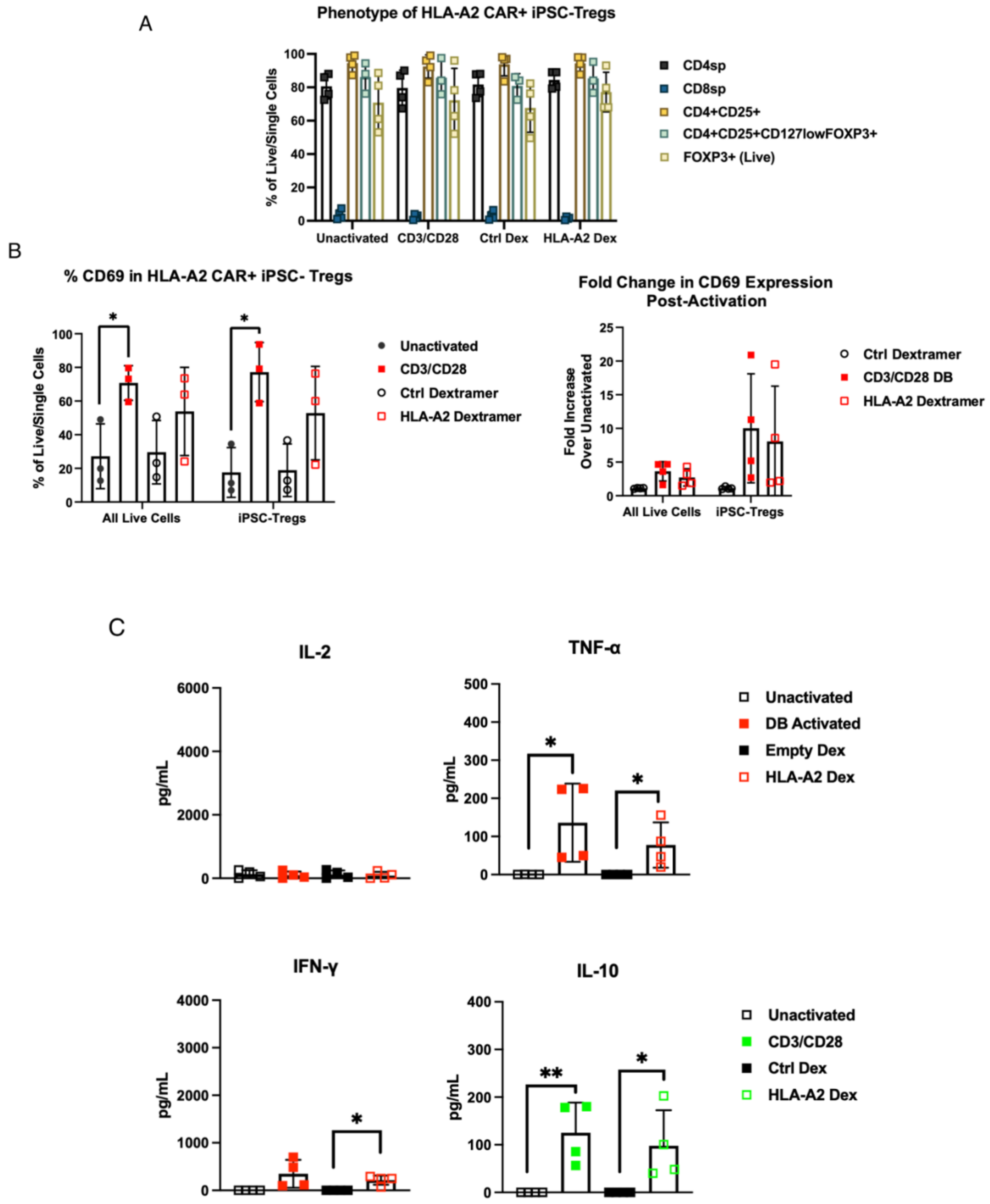

**Supp. Figure 9. Functional characterization of HLA-A2 CAR+ iPSC-Tregs**

A. Expression of Treg markers in sorted unactivated or control dextramer activated HLA-A2 CAR+ iPSC-Tregs and HLA-A2 CAR+ iPSC-Tregs activated via the TCR (CD3/CD28) or HLA-A2 CAR (HLA-A2 dextramer). Phenotype was assessed by flow cytometric analysis (n=4 from n=2 clones).
B. Expression of CD69 in sorted unactivated or control dextramer activated HLA-A2 CAR+ iPSC-Tregs and HLA-A2 CAR+ iPSC-Tregs activated via the TCR or HLA-A2 CAR. The expression of CD69 was assessed in both total live cell populations and target iPSC-Tregs by flow cytometry (left). Fold change in CD69 expression was assessed in activated total live and target iPSC-Tregs compared to unactivated total live cells and target iPSC-Tregs (right, n=4 from n=2 clones).
C. Secretion of cytokines from sorted unactivated or control dextramer activated HLA-A2 CAR+ iPSC-Tregs and HLA-A2 CAR+ iPSC-Tregs activated via the TCR or HLA-A2 CAR (n=4, iPSC-Tregs from n=2 clones).

   Data represent mean ± SD of n independent experiments. $P^* < 0.05$, $P^{**} < 0.01$ by unpaired two-tailed t-test.

**SUPPLEMENTARY FIGURE 10. Characterization of Cryopreserved HLA-A2 CAR+ iPSC-Tregs**

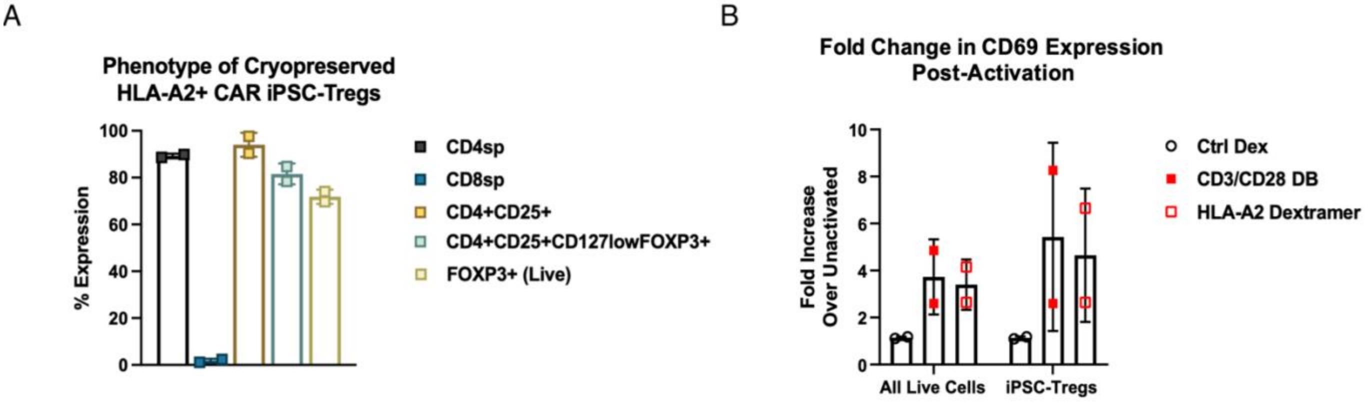


**Supp. Figure 10. Characterization of Cryopreserved HLA-A2 CAR+ iPSC-Tregs**

A. Expression of Treg markers in cryopreserved unactivated HLA-A2 CAR+ iPSC-Tregs. Phenotype was assessed by flow cytometric analysis (n=2 from n=2 clones).
B. Fold increase in CD69 expression was assessed in activated total live and target iPSC-Tregs compared to unactivated total live cells and iPSC-Tregs (n=2 from n=2 clones). Data represent mean ± SD of n independent experiments.
